# Chromatoid body integrates piRNA, SMG6 and m⁶A pathways to control mRNAs in the male germline

**DOI:** 10.64898/2026.03.26.714452

**Authors:** Ammar Ahmedani, Eva Savulkina, Lin Ma, Simang Champramary, Alessia Di Pauli, Elina Valkonen, Samuli OM Laasanen, Riikka Palimo, Ruben Aju George, Keshav Thapa, Tiina Lehtiniemi, Ann-Kristin Dicke, Sabine Kliesch, Nina Neuhaus, Birgit Stallmeyer, Frank Tüttelmann, Juho-Antti Mäkelä, Noora Kotaja

## Abstract

Spermatogenesis requires tightly controlled transcriptome regulation, supported by the PIWI-interacting RNA (piRNA) and nonsense-mediated decay (NMD) pathways, both concentrated in the chromatoid body (CB) of haploid spermatids. We previously showed that the NMD endonuclease SMG6 interacts with the piRNA-binding protein PIWIL1, and that loss of either factor results in overlapping mRNA dysregulation, suggesting functional cooperation. Here, we demonstrate that SMG6 and PIWIL1 assemble with shared RNA-regulatory proteins and bind common mRNA targets in the testis. Functional experiments in GC-2spd cells revealed cooperative regulation of selected transcripts, including *Ccny* and *Taf1d*, and established that SMG6 is required for piRNA-guided degradation of these targets, implicating its endonuclease activity in the piRNA pathway. Target mRNAs were m⁶A-methylated, and this modification shaped their expression levels, SMG6 binding, and CB localization. Moreover, intact target mRNAs accumulated in CBs isolated from *Smg6*-cKO testes, indicating defective CB-associated decay. Together, these findings uncover a key role for the CB in mRNA regulation through coordinated action of the NMD and piRNA pathways.

## INTRODUCTION

Spermatogenesis is a highly specialized differentiation program driven by tightly regulated, stage-specific gene expression networks ^1,2^. Because transcription and translation are uncoupled for extended periods during germ cell differentiation, post-transcriptional regulation becomes essential for maintaining precise temporal control of gene expression and preserving the integrity of the germline transcriptome ^3^. A major component of this regulation involves the formation of ribonucleoprotein (RNP) complexes that coordinate RNA processing, stability, and utilization ^3,4^. RNA quality control and surveillance mechanisms are especially critical during and after meiosis— a phase characterized by broad genome activation and pervasive transcription ^5,6^. The importance of accurate RNA surveillance for normal spermatogenesis and male fertility is underscored by knockout mouse models and by pathogenic mutations in RNA regulatory factors identified in infertile men ^7,8^.

Both germline-specific and broadly conserved RNA regulatory pathways operate in the male germline to support spermatogenesis. Among them, the PIWI-interacting RNA (piRNA) pathway represents a germline-adapted system that evolved to meet the unique demands of germ cell differentiation ^9^. Its best-established function is the silencing of transposable elements, thereby protecting the germline genome from transposon activation and invasion ^10^. Beyond this, piRNAs are also implicated in regulating coding and non-coding RNAs, although these roles remain incompletely defined ^11^. piRNA functions are mediated by PIWI proteins: PIWIL2/MILI acts from early germ cell stages through early haploid round spermatids, while PIWIL1/MIWI predominates in round spermatids ^12,13^. Complementing germline-specific mechanisms, conserved RNA quality-control pathways such as nonsense-mediated mRNA decay (NMD) ensures transcriptome fidelity by selectively degrading mRNAs with features like premature termination codons or long 3′ UTRs ^14^. Genetic disruption of key NMD factors such as the effectors UPF2 and UPF3A or the endonuclease SMG6 demonstrates that NMD is essential for normal spermatogenesis and male fertility ^15–17^.

Interestingly, the components of both the piRNA and NMD pathways accumulate in the chromatoid body (CB), a phase-separated cytoplasmic RNP granule in haploid round spermatids ^17–19^. The CB is an unusually large, germline-specific RNP granule (germ granule) ^20,21^. Its precursors appear in late pachytene spermatocytes and later coalesce into a single perinuclear granule after meiosis ^21^. Our previous work demonstrated that the CB is enriched with RNA-binding proteins that recruit diverse RNAs, including piRNAs, mRNAs, and intergenic non-coding transcripts, highlighting its role as an organizational hub for RNA regulation during spermiogenesis ^18^.

Despite increasing insight into the CB composition, the mechanisms that target specific RNAs to this structure remain largely unresolved. Intriguingly, the CB has been reported to enrich several N6-methyladenosine (m⁶A) reader proteins, including IGF2BP3, YTHDC2 and PRRC2A ^18,22,23^. The m⁶A modification is written by the METTL3-METTL14 complex, erased by demethylases (FTO, ALKBH5), and recognized by reader proteins to control post-transcriptional RNA metabolism, including stability, translation, subcellular localization and decay ^24–26^. m⁶A writers, erasers and readers have emerged as critical regulators of germ cell differentiation ^27^. The selective accumulation of multiple m⁶A readers within the CB therefore raises the intriguing possibility that m⁶A marks contribute to the targeting and/or retention of RNAs in this germline-specific RNP granule.

Previously, we showed that the NMD endonuclease SMG6 and the piRNA-binding protein PIWIL1 are highly enriched in the CB and exhibit nearly identical spatiotemporal expression patterns, both proteins first appearing in CB precursors and persisting within the CB throughout round spermatid differentiation ^17^. Knockout mouse model revealed that SMG6 is essential for transcriptome regulation in male germ cells ^17^. Its deletion disrupts spermatogenesis, causing aberrant mRNA accumulation, particularly the transcripts with long 3′UTRs. *Piwil1* deletion produces a strikingly similar phenotype and dysregulation of a shared set of mRNAs ^17,28^. We further showed that PIWIL1 and SMG6 physically interact in the testis, suggesting functional cooperation between the piRNA pathway and NMD in coordinating mRNA regulation within the CB ^17^. However, despite this strong phenotypic, molecular, and spatial convergence, the mechanistic basis of how these two pathways intersect and are coordinated within the CB remains largely unresolved.

In this study, we combine biochemical, imaging, and functional analyses in GC-2spd cells, mouse models, and human testis samples to uncover the mechanisms underlying CB-localized cooperation between the piRNA and NMD pathways. Our findings reveal a functional integration of piRNA-mediated regulation with SMG6 endonuclease activity and identify m⁶A as a key signal directing mRNAs to the CB and to SMG6-dependent decay. Together, these results establish the CB as a spatiotemporal hub for selective transcript turnover coordinated by PIWIL1, SMG6, and m⁶A-dependent RNA recognition.

## RESULTS

### SMG6 and PIWIL1 form RNA regulatory complexes in the testis

To explore the potential cooperation of SMG6 and PIWIL1 in male germ cells, we first identified SMG6- and PIWIL1-interacting proteins in the adult mouse testis by immunoprecipitation (IP) followed by mass spectrometry (Fig. 1A). We included only the hits with at least 2 peptides in all three replicates but not present in the IgG control samples. We identified 717 SMG6-interacting proteins and 569 PIWIL1-interacting proteins (Supplementary Table S1A,B). Notably, 470 proteins co-precipitated with both SMG6 and PIWIL1, indicating a substantial overlap and suggesting that they are part of overlapping ribonucleoprotein complexes (Fig. 1B, Supplementary Table S1A,B). Notably, SMG6 and PIWIL1 were both found in reciprocal experiments, supporting a specific interaction between the two proteins. 36 of the established CB proteins (>40%) ^18^ were detected among the shared interaction partners (Fig. 1B, Supplementary Table S1A,B), suggesting that many of the interactions occur within the CB compartment. Functional enrichment analysis of the shared interaction partners revealed significant enrichment of pathways associated with post-transcriptional gene regulation, including translation, NMD, splicing, and RNP binding and RNA localization. (Fig. 1C, Supplementary Table S1C). Interestingly, both SMG6 and PIWIL1 also co-purified with several m⁶A readers, including PRRC2A, YTHDC2, and HNRNPC, linking their function to m⁶A-modified RNA regulation.

**Figure 1.**
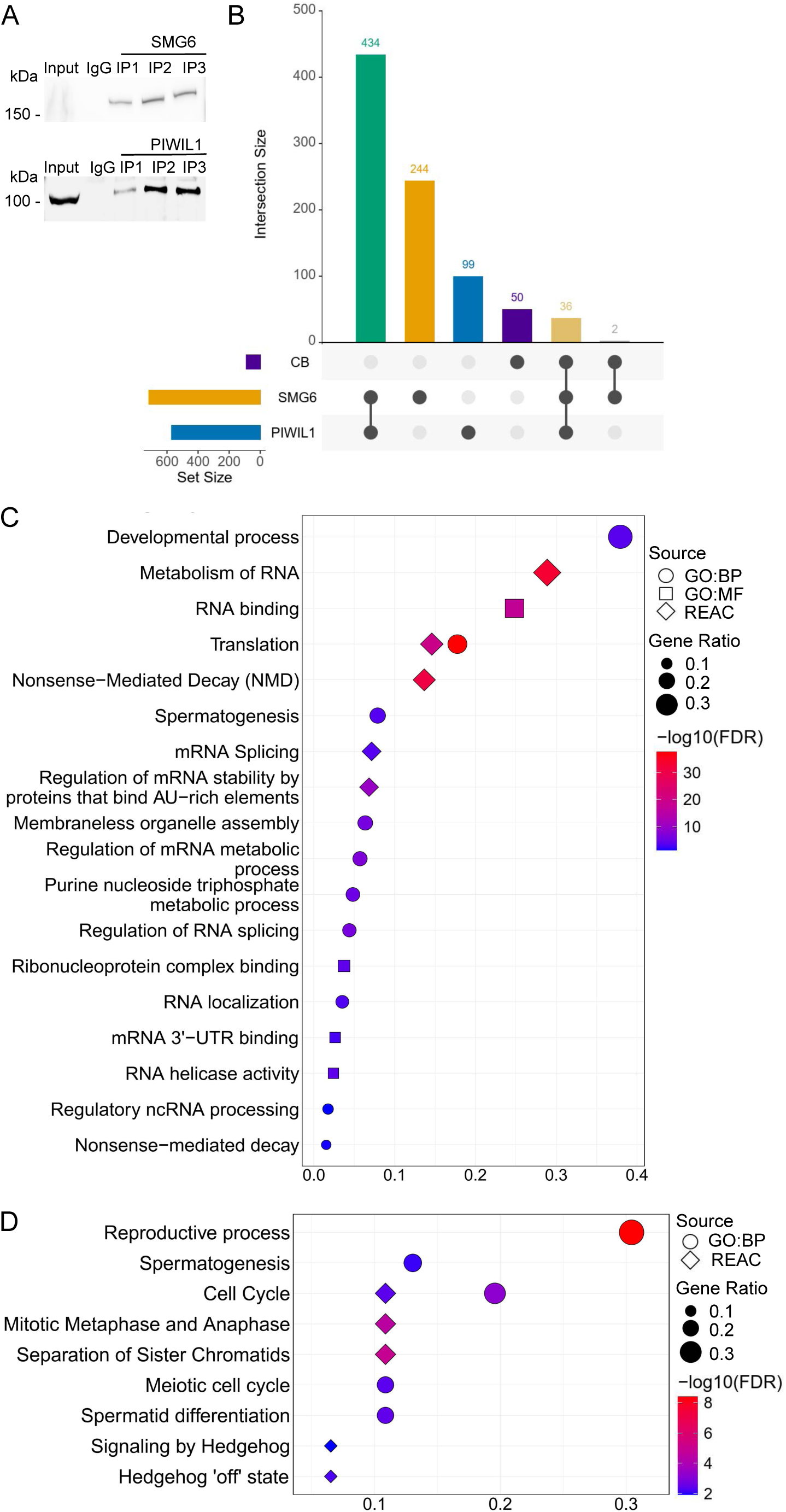
SMG6 and PIWIL1 associate with overlapping set of proteins and mRNAs in mouse testes. A) Validation of the anti-SMG6 and anti-PIWIL1 IP samples submitted to mass spectrometry by immunoblotting with the same antibodies. IP was done from adult mouse testes. Rabbit IgG was used as a negative control. B) UpSet plot showing the intersections among proteins identified in PIWIL1, SMG6, and CB-associated mass spectrometry datasets. Vertical bars indicate the number of proteins shared between the indicated dataset combinations, with connected dots below each bar denoting the corresponding intersection. Horizontal bars represent the total number of proteins detected in each dataset. Numbers above the bars indicate intersection sizes. C) Functional enrichment analysis of shared SMG6 and PIWIL1 interaction partners. Dot plot of enriched Gene Ontology (GO) terms and Reactome pathways identified by g:Profiler, highlighting biologically relevant, significantly enriched categories. The x-axis shows the gene ratio and the y-axis the corresponding term descriptions. Color intensity reflects statistical significance (−log10(FDR); FDR < 0.05) with a blue-to-red gradient, where red denotes higher significance. Dot shape indicates the annotation source: circles, GO Biological Process (GO:BP); squares, GO Molecular Function (GO:MF); triangles, Reactome pathways. D) Functional enrichment analysis of shared SMG6 and PIWIL1 bound mRNAs.

In addition to sharing protein interaction partners, IP followed by RNA sequencing (RIP-seq) revealed that SMG6 and PIWIL1 also form complexes with a similar set of mRNAs (Supplementary Table S2A,B). We sequenced a total of three anti-SMG6, three anti-PIWIL1, and three control IgG samples immunoprecipitated from adult mouse testes (Supplementary Fig. S1A). Comparison of the IP samples to the IgG controls revealed 156 mRNAs specifically enriched in the SMG6 complexes and 182 mRNAs enriched in the PIWIL1 complexes (Supplementary Table S2A,B). Strikingly, 49 of these RNAs were found in both the SMG6 and PIWIL1 complexes (Supplementary Fig. S1B, Supplementary Table S2A,B). Functional enrichment analysis revealed that these shared mRNAs encoded for genes involved in important spermatogenic processes, including meiosis (Fig. 1D, Supplementary Table S2C). Selective enrichment of mRNAs encoding for meiotic regulators, such as *Pttg1* (inhibits chromosome segregation), *Cks2* (required for metaphase to anaphase transition), *Xlr5c* (meiotic chromosome/synaptonemal complex protein), and *Ccnb1ip1* (ubiquitin E3 ligase for crossover formation), supports the previously suggested role for the SMG6-PIWIL1 complex in the regulation of the meiotic-to-post-meiotic gene expression program ^1^. Taken together, these data indicate that SMG6 and PIWIL1 form interconnected complexes in the testis that enable their cooperation in post-transcriptional mRNA regulation.

### SMG6 and PIWIL1 share target mRNAs in the testis

Next, we aimed to identify potential target mRNAs whose stability is affected by both SMG6 and PIWIL1 *in vivo*. We first revisited our existing mRNA-seq data ^17^, which revealed that ∼300 genes were upregulated in both *Smg6-cKO* (GSE182518) and *Piwil1-KO* (GSE64138) round spermatids (Log2FC≥1, Padj≤0.05) (Supplementary Table S3A). From this set, we then selected transcripts with a reduced number of degradation fragments in *Smg6*-cKO, which is indicative of direct endonucleolytic targeting. To this end, we performed degradome sequencing (degradome-seq) for *Smg6*-cKO and control round spermatids (3 biological replicates each). We focused our analysis of the degradome-seq data only on the 4812 genes that were found to be upregulated in *Smg6*-cKO vs. control round spermatids via RNA-seq ^17^ to identify the upregulated genes whose mRNA degradation products decreased in the absence of SMG6. After filtering out the reads with low expression, we identified degradation products from 140 of these upregulated genes. Strikingly, DE analysis revealed that 77 of these genes had significantly downregulated degradation products (altogether 126 downregulated degradation products) in *Smg6*-cKO round spermatids, with only 3 genes exhibiting significantly upregulated degradation products (Supplementary Table S3B, Fig. 2A). Thirteen of the 77 genes with decreased degradation products were found to be among the ∼300 shared targets, *i.e.* they were also upregulated in *Piwil1*-KO round spermatids (Supplementary Table S3B; *Taf1d, Iqcb1, Hnrnpul2, 4833439L19Rik, Tpcn2, Skap2, Ccny, Hspa8, Sf3b6, Ppp1r37, Zmym1, Tfrc, Man2a2*). These 13 genes that are upregulated in both *Smg6*-cKO and *Piwil1*-KO round spermatids and exhibit reduced degradation intermediates in the absence of SMG6 are likely targets for SMG6-PIWIL1-mediated degradation.

**Figure 2.**
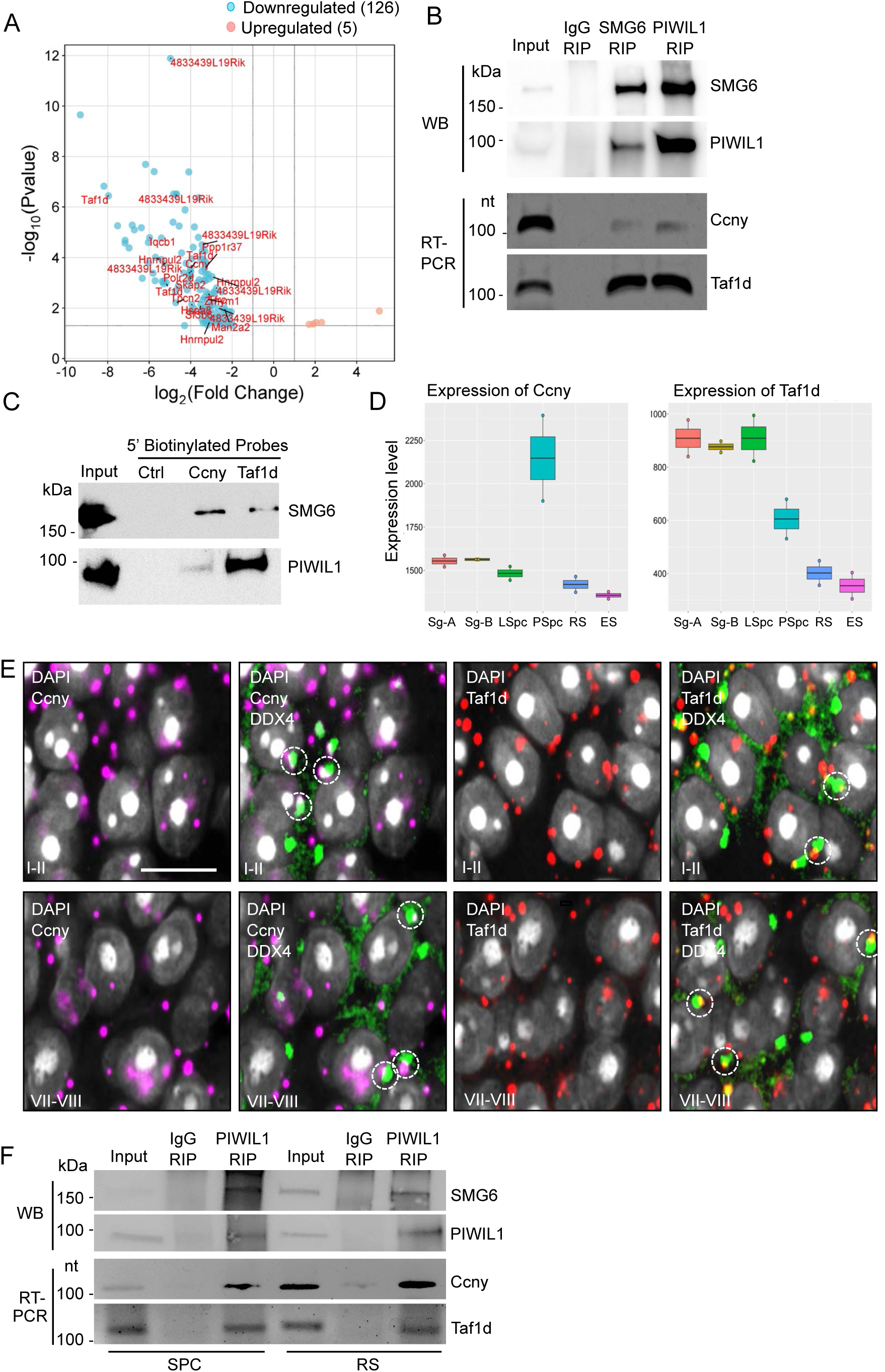
SMG6 and PIWIL1 associate with *Ccny* and *Taf1d* mRNAs in mouse testes. A) Volcano plot shows the downregulated degradation products for the shared upregulated genes in *Piwil1*-KO and *Smg6*-cKO round spermatids. B) SMG6 and PIWIL1 form complexes with each other and with *Ccny* and *Taf1d* mRNA in mouse testes. Upper panels show Western blotting (WB) of SMG6 and PIWIL1. Lower panels show RT-PCR using *Ccny* and *Taf1d* specific primers. C) RNA pulldown assay from adult mouse testicular lysate using biotinylated DNA probes complementary to *Ccny* and *Taf1d* transcripts and streptavidin beads, followed by Western blotting with anti-SMG6 and anti-PIWIL1 antibodies. Scrambled probe was used as a negative control (Ctrl). D) Expression of *Ccny* (left) and *Taf1d* (right) in type A spermatogonia (Sg-A), type B spermatogonia (Sg-B), leptotene spermatocytes (LSpc), pachytene spermatocytes (PSpc), round spermatids (RS) and elongating spermatids (ES) (GSE35005). E) *In situ* hybridization of mouse testis sections using probes detecting *Ccny* (magenta) and *Taf1d* (red). CB was labeled with anti-DDX4 antibody (green). DAPI stains the nuclei (grey). Example stainings of the stages I-II and VII-VIII of the seminiferous epithelial cycle are shown. Selected *Ccny-* and *Taf1d-*associated CBs are circled. Scale bar: 10 µm. F) Upper panels show Western blotting (WB) of PIWIL1 IP from spermatocytes (SPC) and round spermatids (RS) using anti-PIWIL1 and anti-SMG6 antibodies. Lower panels show RT-PCR after IP using *Ccny* and *Taf1d* specific primers.

### Shared target mRNAs *Ccny* and *Taf1d* associate with SMG6 and PIWIL1 in the testis

Among the identified shared targets of SMG6 and PIWIL1, *Ccny* and *Taf1d* were selected for further functional studies. TAF1D/TAF41 is a subunit of the SL1/TIF-IB complex that is required for RNA polymerase I pre-initiation complex assembly and rDNA promoter recognition ^29^, while CCNY is a conserved cyclin family protein that forms active complexes with CDK14 and CDK16, regulating kinase activity and diverse cellular processes ^30^. IP of SMG6 and PIWIL1 from adult mouse testes, followed by RT-PCR, validated the associations of *Ccny* and *Taf1d* with SMG6 and PIWIL1 in the testis (Fig 2B). To further validate the physical interaction between the PIWIL1–SMG6 complex and its mRNA substrates, we performed RNA pulldown assays using biotinylated DNA probes complementary to the *Ccny* and *Taf1d* transcripts. Streptavidin-based pulldown followed by Western blotting revealed that both SMG6 and PIWIL1 were efficiently recovered from testis lysates with *Ccny* and *Taf1d*-specific probes (Fig. 2C). These reciprocal approaches confirm the association of *Ccny* and *Taf1d* with PIWIL1 and SMG6, suggesting that they can be targeted by the PIWIL1/SMG6 complex in the testis.

To pinpoint the phase of spermatogenesis when SMG6/PIWIL1-mediated regulation of *Ccny* and *Taf1d* could occur, we first studied their expression profile in differentiating male germ cells by existing mRNA-seq data (GSE35005) ^2^. *Ccny* expression was shown to peak in pachytene spermatocytes and it was quickly downregulated in haploid spermatids (Fig. 2D). *Taf1d* was expressed at the highest levels during early phase of spermatogenesis, followed by a progressive decline in post-meiotic cells (Fig. 2D). Fluorescence *in situ* hybridization confirmed a strong expression of both *Ccny* and *Taf1d* in spermatocytes and the gradual decrease of their expression during spermatid differentiation (Supplementary Fig. S2). Combining the *in situ* hybridization signal with DDX4 immunofluorescence (IF) revealed the association of both the *Ccny* and *Taf1d* signals with the CB, confirming their spatial localization to this RNP granule (Fig. 2E). The expression of *Ccny* and *Taf1d* coincided with the expression of SMG6 and PIWIL1, which is induced at the time of the formation of the CB during the transition from late spermatocytes to round spermatids ^17^. IP of PIWIL1 from isolated spermatocytes and round spermatids confirmed that PIWIL1 forms complexes with SMG6, *Ccny* and *Taf1d* in both cell types (Fig. 2F). In conclusion, the expression and localization patterns of *Ccny* and *Taf1d* support the role of CB-localized SMG6-PIWIL1 in their regulation in late spermatocytes and round spermatids.

### PIWIL1 facilitates SMG6-mediated mRNA decay in GC-2spd cells

To further investigate the functional cooperation between SMG6 and PIWIL1, we utilized GC-2spd cells, a mouse spermatogenic cell line derived from pachytene spermatocytes, as a model ^31,32^. These cells represent a well-established *in vitro* model for studying germ cell-related processes, and they endogenously express SMG6, PIWIL1, and piRNAs, making them a suitable system for dissecting the interplay between the NMD endonuclease SMG6 and the piRNA-binding PIWIL1. We first performed co-IP of SMG6 and PIWIL1 from GC-2spd cells, followed by Western blotting, and showed that PIWIL1 and SMG6 interact in cultured germ cells even without the CB environment (Fig. 3A). Furthermore, RT-PCR after IP revealed that these proteins also form complexes with the *Ccny* and *Taf1d* mRNAs in cultured cells (Fig. 3A).

**Figure 3.**
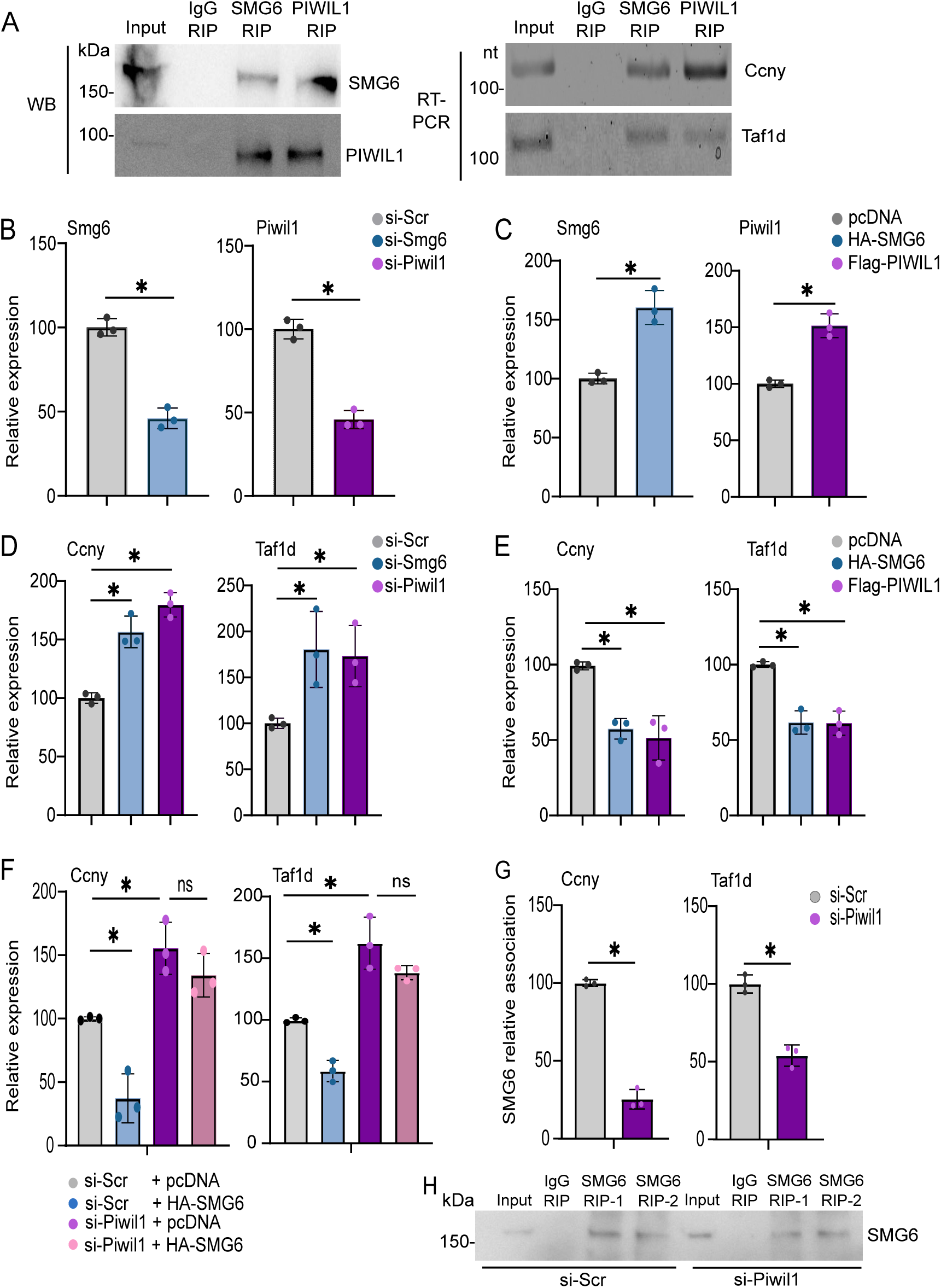
SMG6 and PIWIL1 regulate *Taf1d* and *Ccny* expression in GC-2spd cells. A) RNA immunoprecipitation (RIP) from GC-2spc cells using SMG6 and PIWIL1 antibodies. Rabbit IgG was used as a negative control. Left panel shows Western blotting (WB) of SMG6 and PIWIL1. Right panel shows RT-PCR after IP using *Ccny* and *Taf1d* specific primers. B) Validation of the knockdown efficiency after transfection of small interfering RNA (siRNA) for *Smg6* or *Piwil1* (si-Smg6, si-Piwil1) in GC-2spd cells by RT-qPCR using *Smg6* or *Piwil1* primers. Scrambled siRNA was used as a negative control (si-Scr). C) Validation of the overexpression of SMG6 or PIWIL1 in GC-2spd cells using HA-SMG6 or FLAG-PIWIL1 expression plasmids, respectively. Empty pcdna3.1 vector as a control. D) RT-qPCR using *Ccny* and *Taf1d*-specific primers after siRNA-mediated downregulation of *Smg6* or *Piwil1* in GC-2spd cells. E) RT-qPCR using *Ccny* and *Taf1d*-specific primers after overexpression of HA-SMG6 or FLAG-PIWIL1 in GC-2spd cells. F) Overexpression of SMG6 (HA-SMG6) in GC-2spd cells in the presence or absence of *Piwil1* knockdown by siRNA. RT-qPCR analysis of *Ccny* and *Taf1d* mRNAs was performed to assess whether PIWIL1 is required for SMG6-mediated regulation of these targets. G) Control (si-Scr) or PIWIL1 knockdown (si-Piwil1) GC-2spd cells were subjected to SMG6 RIP, followed by RT-qPCR to assess the association of SMG6 with *Ccny* and *Taf1d* mRNAs. *Taf1d* and *Ccny* transcripts were quantified and normalized to input RNA. Data represent mean ± S.D. from three independent experiments, *P < 0.05. H) Western blotting with anti-SMG6 antibody to validate equal IP of SMG6 from control (si-Scr) and *Piwil1*-silenced (si- Piwil1) cells. Two representative IPs (RIP-1, RIP-2) are shown for both conditions. For all graphs (B-F), means of three independent experiments are shown, and *Rpl19* was used as a reference gene. Data represent mean ± S.D. from three independent experiments, *P < 0.05.

A targeted knockdown of *Smg6* or *Piwil1* efficiently reduced their mRNA levels, with siRNAs decreasing their expression to ∼50% of control (Fig. 3B), while ectopic expression of HA-SMG6 or Flag-PIWIL1 elevated their own mRNA levels to ∼150% (Fig. 3C). Importantly, siRNA-mediated depletion of either *Smg6* or *Piwil1* led to a pronounced upregulation of *Ccny* and *Taf1d* (Fig. 3D). This finding suggests that the upregulation of *Ccny* and *Taf1d* in *Smg6*-cKO and *Piwil1*-KO round spermatids *in vivo* is likely due to the direct role of SMG6 and PIWIL1 in their regulation. Conversely, overexpression of either factor resulted in robust transcript downregulation (Fig. 3E). Notably, SMG6 overexpression failed to suppress *Ccny* and *Taf1d* levels in PIWIL1-depleted cells (Fig. 3F), indicating that PIWIL1 is required for SMG6-mediated mRNA decay. Furthermore, PIWIL1 knockdown significantly reduced *Ccny* and *Taf1d* mRNA association with SMG6 as shown by SMG6-RIP followed by RT-PCR (Fig. 3G,H). These results suggest that PIWIL1 facilitates the association of SMG6 with its target mRNAs, thereby coupling PIWIL1 with SMG6-mediated endonucleolytic mRNA decay in germline-derived cells.

### SMG6 and PIWIL1 regulate m⁶A-modified *Ccny* and *Taf1d* in GC-2spd cells

The identification of m⁶A readers as SMG6/PIWIL1-interacting proteins in the testis (Supplementary Table S1) prompted us to study whether m⁶A modification is involved in SMG6/PIWIL1-mediated mRNA regulation. m⁶A IP followed by RT–PCR revealed robust enrichment of both *Ccny* and *Taf1d* in the m⁶A-modified RNA fraction, confirming that they are m⁶A-modified in GC-2spd cells (Fig. 4A). To test whether m⁶A modification contributes to the control of their expression level, we inhibited methylation through combined genetic and pharmacological approaches. siRNA-mediated knockdown of *Mettl3* significantly reduced METTL3 protein levels and resulted in accumulation of *Ccny* and *Taf1d* mRNAs compared to scramble controls (Fig. 4B,C). Furthermore, treatment with the methyltransferase inhibitor STC-15 led to a clear accumulation of *Ccny* and *Taf1d* transcripts, as revealed by both RT-PCR and *in situ* hybridization, accompanied by a reduction in global m⁶A levels as confirmed by m⁶A dot blot analysis (Fig. 4D,E, Supplementary Fig. S3). These findings suggest that m⁶A methylation promotes the degradation of *Ccny* and *Taf1d* mRNAs.

**Figure 4.**
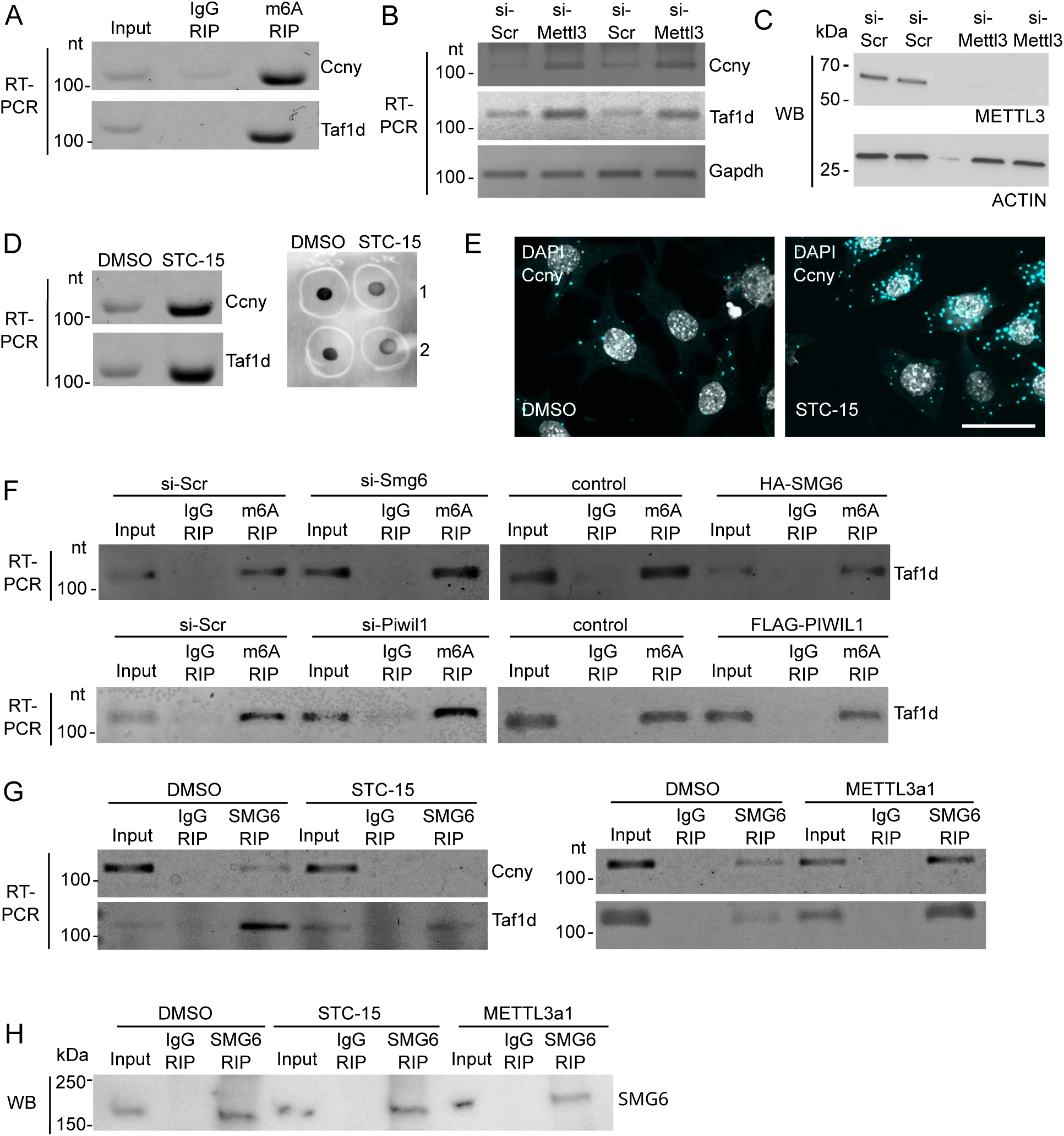
SMG6 and PIWIL1 promote decay of m⁶A-modified *Ccny* and *Taf1d* in GC-2spd cells. A) m⁶A -RIP followed by RT-PCR using primers for *Taf1d* and *Ccny.* Mouse IgG was used as a control. B) METTL3 knockdown by siRNA (si-Mettl3) followed by RT-PCR to amplify *Ccny* and *Taf1d* mRNAs. Scrambled siRNA (si-Scr) was used as a control. C) Downregulation of METTL3 was validated by anti-METTL3 western blotting. Antibody against ACTIN was used as a loading control. D) GC-2spd cells were treated with the methyltransferase inhibitor STC-15 followed by RT-PCR using *Ccny* and *Taf1d* primers (left panel). Right panel shows the levels of m⁶A RNA in control (DMSO) and STC-15-treated cells by dot blotting with anti-m6A antibody (two independent replicates are shown). E) *In situ* hybridization using *Ccny*-specific probe of DMSO- or STC-15 treated GC-2spd cells. DAPI stains the nuclei (grey). Scale bar: 10 µm. F) GC-2spd cells were transfected with control siRNA (si-Scr), *Smg6*- or *Piwil1*-targeting siRNA (si-Smg6 or si-Piwil1), empty pcdna3.1 plasmid (control) or HA-SMG6 and FLAG-PIWIL1 overexpressing plasmids before m⁶A - RIP. m6-A-modified *Taf1d* was detected with RT-PCR from anti-m⁶A RIP samples. Mouse IgG was used as a control in the RIP. G) GC-2spd cells were treated with DMSO, METTL3a1, or STC-15 before anti-SMG6-RIP. SMG6-associated *Ccny* and *Taf1d* mRNAs were detected with RT-PCR. Rabbit IgG was used as a control in the RIP. H) Western blot with anti-SMG6 antibody to validate equal IP of SMG6 from DMSO, STC-15, and METTL3a1 treated cells. One representative RIP experiment is shown for each condition.

Guided by this observation, we next examined the interconnection between the m⁶A modification and SMG6/PIWIL1 function. To this end, we first studied whether the modulation of SMG6 and PIWIL1 expression affects the levels of m⁶A-modified *Ccny* and *Taf1d* by m⁶A-IP, followed by RT–PCR. Interestingly, depletion of SMG6 or PIWIL1 led to a marked accumulation of m⁶A-modified *Taf1d* mRNAs, whereas overexpression of either protein caused a reduction of these methylated species (Fig. 4F), suggesting that the SMG6/PIWIL1 complex regulates the methylated fractions of *Taf1d* mRNAs in GC-2spd cells. Next, we investigated whether changes in m⁶A deposition influence the association of SMG6 endonuclease with its target mRNAs by modulating m⁶A levels prior to SMG6-RIP. We showed that reducing m⁶A levels by STC-15 treatment markedly diminished SMG6 binding to *Ccny* and *Taf1d* mRNAs compared with that in the Dimethyl sulfoxide (DMSO) controls (Fig. 4G,H). Conversely, enhancing methylation with METTL3 activator METTL3a1 increased SMG6 binding to both mRNAs (Fig. 4G,H). Together, these results support the role of m⁶A deposition in promoting the recruitment of target RNAs to the PIWIL1–SMG6 complex for degradation.

### m⁶A modification and SMG6 are required for piRNA-induced mRNA decay in GC-2spd cells

PIWIL1 and other PIWI proteins bind piRNAs to mediate their functions; however, PIWI proteins are also known to have piRNA-independent functions ^33^. To study if piRNAs are involved in the SMG6-PIWIL1-mediated mRNA decay, we first showed that piRNAs are present in the PIWIL1/SMG6 complexes immunoprecipitated from adult mouse testes (Fig. 5A). To test whether piRNAs can induce the degradation of transcripts co-regulated by SMG6 and PIWIL1, we transfected GC-2spd cells with RNAs immunoprecipitated with PIWIL1 from adult mouse testes, and showed that transfection of PIWIL1-associated RNAs induced the downregulation of *Ccny* mRNA (Fig. 5B).

**Figure 5.**
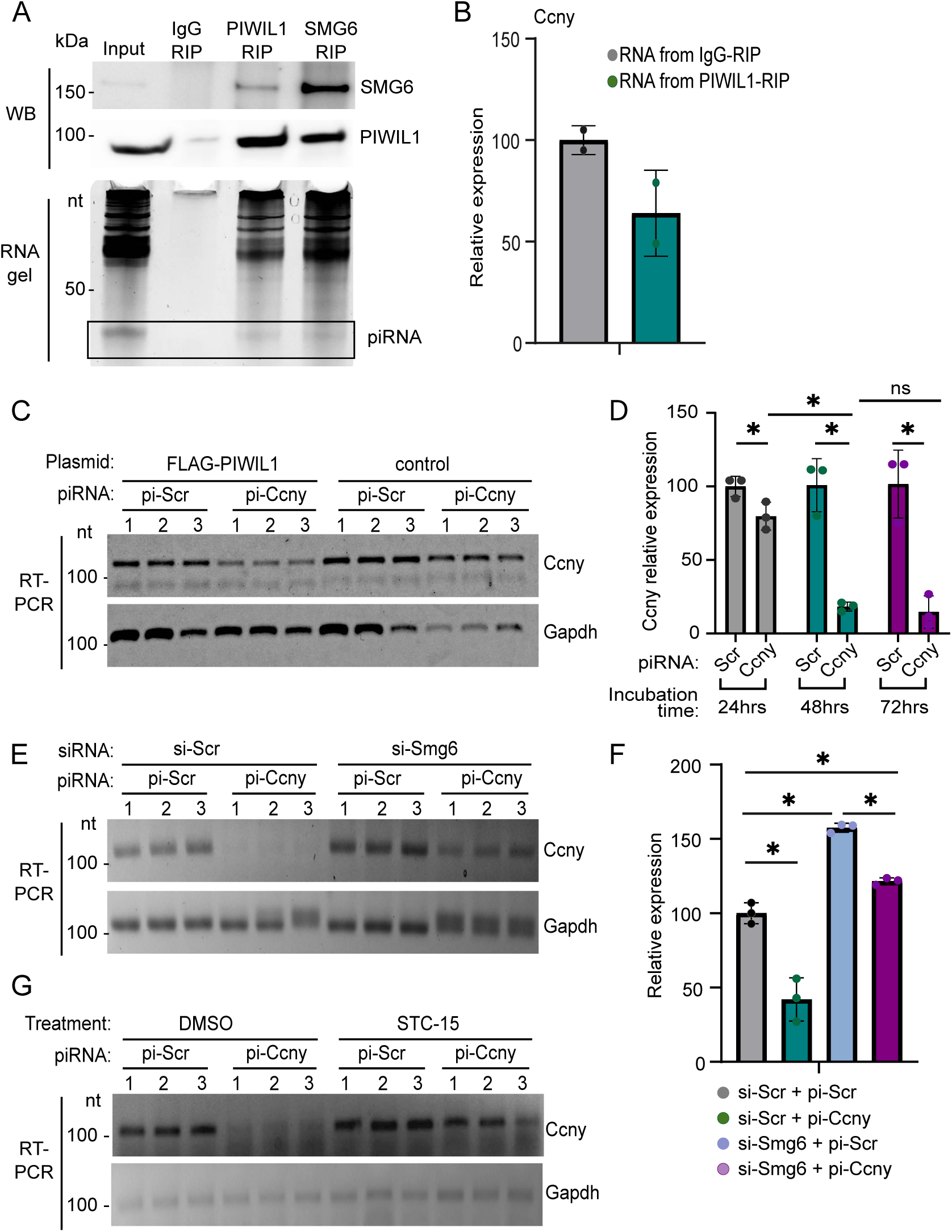
m⁶A modification and SMG6 are required for piRNA-induced mRNA decay. A) Immunoprecipitation of PIWIL1 and SMG6 from adult mouse testes. Successful IP was confirmed with Western blotting (WB) using SMG6 and PIWIL1 antibodies. Associated small RNAs were visualized in the UREA-PAGE gel with SYBR Gold staining, revealing a 26-32-nt population consistent with piRNAs length. B) Transfection of PIWIL1-associated testis RNAs into GC2spd cells, followed by RT-qPCR analysis of *Ccny* target mRNA. *Gapdh* was used as a reference gene. Data represent mean ± S.D. from two independent experiments. C) Transfection of *Ccny*-targeting piRNA (*pi-Ccny*) induces *Ccny* mRNA downregulation in 3T3 cells only when PIWIL1 is overexpressed. Scrambled piRNA (*pi-Scr*) was used as a control for *Ccny*-targeting piRNA. Empty pcdna3.1. vector (control) was used as a control for FLAG-PIWIL1 overexpression. D) GC-2spd cells were transfected with *pi-Scr* or *pi-Ccny*, followed by RT-qPCR using *Ccny* primers at different time points after transfection to monitor *Ccny* mRNA levels. E) GC-2spd cells were subjected to siRNA-mediated knockdown of *Smg6* prior to transfection with the synthetic *pi-Ccny*, followed by RT-PCR analysis of *Ccny* mRNA. F) Same experiment as in (E), but *Ccny* expression was detected by RT-qPCR. G) GC-2spd cells were treated with the METTL3 inhibitor STC-15 to block m⁶A deposition, followed by transfection with the synthetic *pi-Ccny* and RT-PCR analysis of *Ccny* mRNA. For graphs (D,F), *Gapdh* was used as a reference gene. Data represent mean ± S.D. from three independent experiments; *P < 0.05.

We next focused on specific piRNAs targeting *Ccny* mRNA. The screening of the piRNA population expressed in round spermatids using our existing small-RNAseq data ^34^ identified a candidate with strong complementarity to the *Ccny* 3′UTR (Supplementary Fig. S3B). We first transfected this *Ccny*targeting synthetic piRNA (pi-*Ccny*) into somatic 3T3 cells, a mouse embryonic fibroblast line that does not express endogenous PIWI proteins or piRNAs. In these cells, neither pi-*Ccny* nor overexpressed PIWIL1 alone were able to downregulate *Ccny* levels. However, transfection of pi-*Ccny* with simultaneous overexpression of PIWIL1 induced *Ccny* mRNA downregulation (Fig. 5C), indicating that both piRNA and PIWIL1 are needed to bring about the effect. In line with this, transfection of pi-*Ccny* into GC-2spd cells, which endogenously express PIWIL1, resulted in a marked reduction of *Ccny* mRNA levels (Fig. 5D). Notably, SMG6 knockdown substantially hampered the piRNA-mediated effect (Fig. 5E,F), revealing that SMG6 is required for piRNA-induced *Ccny* downregulation.

Building on our previous findings that m⁶A contributes to SMG6/PIWIL1-mediated regulation, we also tested whether m⁶A is necessary for piRNA-guided SMG6-dependent decay. Interestingly, inhibition of m⁶A deposition via the METTL3 inhibitor STC-15 efficiently blocked the piRNA-mediated *Ccny* decay (Fig. 5G). Altogether, these results demonstrate that m⁶A modification and the NMD endonuclease SMG6 are both required for piRNA-induced mRNA decay in GC-2spd cells, revealing m⁶A as a licensing signal for PIWIL1/piRNA-mediated transcript cleavage and SMG6 as a novel downstream effector for piRNA action.

### m⁶A modified RNAs associate with SMG6 and PIWIL1 in the testis and are enriched in the CB

After confirming the functional cooperation of PIWIL1 and SMG6 in cultured cells, as well as the involvement of m⁶A modification in the PIWIL1/SMG6-mediated mRNA decay, we wanted to translate these findings to the *in vivo* environment in the testis. To study whether PIWIL1 and SMG6 associate with m⁶A -modified RNAs in the testis, we performed IP using an anti-m^6^A antibody to pull down all m⁶A-modified RNAs. Western blotting of the immunoprecipitated samples revealed that both PIWIL1 and SMG6 co-purify with m⁶A-modified RNAs in the adult mouse testis (Fig 6A). Conversely, IP of PIWIL1 and SMG6 recovered m⁶A -modified transcripts, as shown by dot blotting with an m⁶A antibody (Fig. 6B). These findings reinforce the idea that SMG6 and PIWIL1 interact with m⁶A-modified transcripts, potentially also regulating their fate in germ cells *in vivo*.

**Figure 6.**
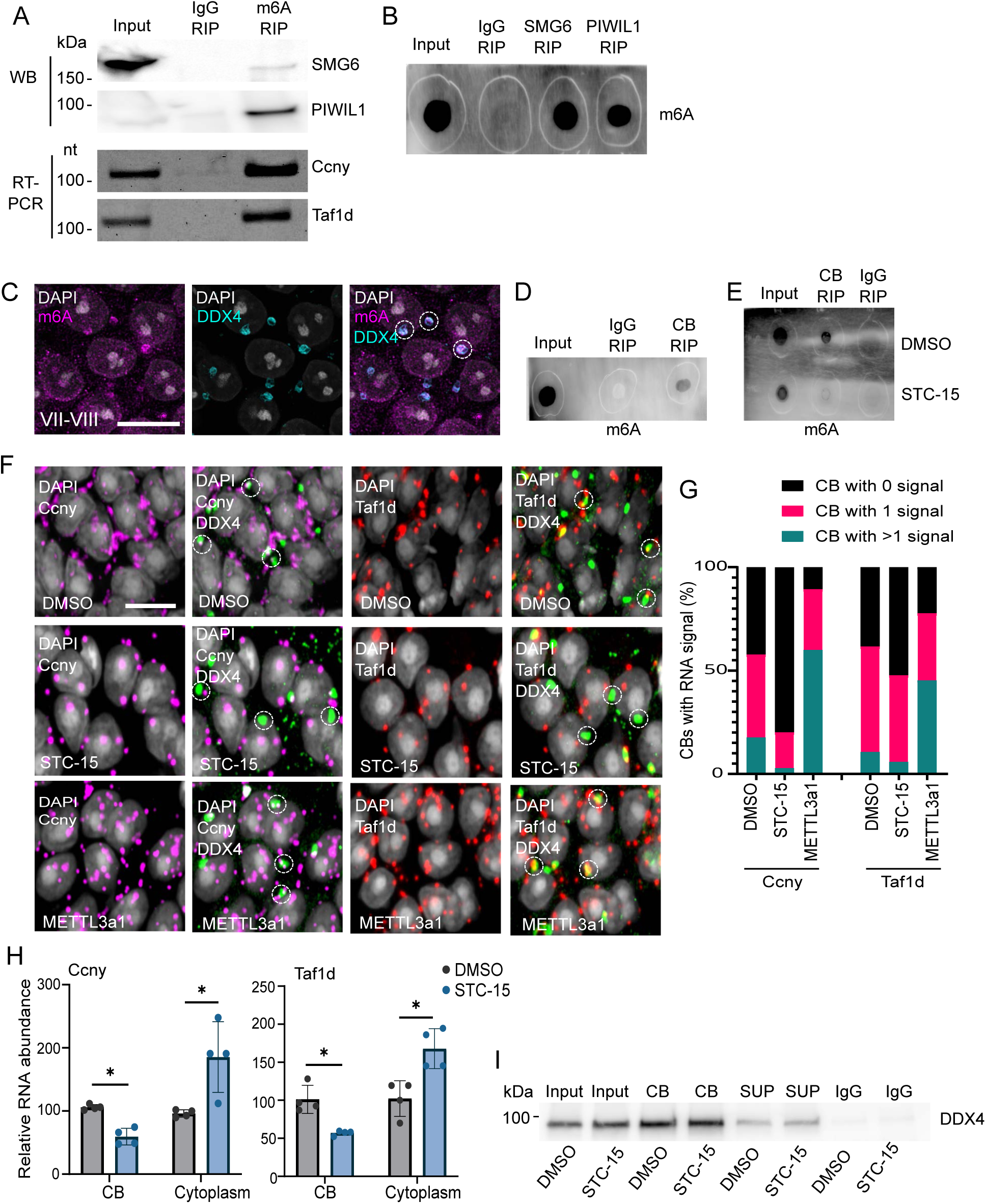
SMG6, PIWIL1 and the CB are associated with m⁶A -modified RNAs in the testis. A) m⁶A RIP from adult mouse testis lysates followed by Western blotting (WB) with anti-SMG6 and anti-PIWIL1 antibodies, or RT-PCR using primers for *Ccny* and *Taf1d*. Mouse IgG was used as a negative control in the RIP. B) RIP using anti-SMG6 or anti-PIWIL1 from testis lysates followed by dot blotting with anti-m⁶A antibody shows the enrichment of m⁶A -modified RNAs in the SMG6 and PIWIL1 complexes. C) IF staining of PFA-fixed paraffin-embedded adult mouse testis section (stage VII-VIII) with anti-m⁶A antibody (magenta). Anti-DDX4 antibody was used to label the CB (cyan). DAPI stains the nuclei. Scale bar: 10 µm D) Dot blotting of RNA extracted from CBs isolated from adult mouse testes (CB-RIP) with anti-m⁶A antibody. E) Dot blotting of CBs isolated from DMSO or STC-15 treated cultured testicular cells. F) *In situ* hybridization of squash preparations after *ex vivo* culture of the tubule fragments (stage II-V) in the presence of STC-15 or METTL3a1 using probes detecting *Ccny* (magenta) or *Taf1d* (red). DMSO-treated tubules were used as a control. CB were labeled with anti-DDX4 antibody (green). DAPI stains the nuclei (grey). Scale bar: 10 µm. G) Quantification of the co-localization of *Taf1d* and *Ccny* mRNA-foci with CBs. Data represent the percentage of CBs with no overlapping *Ccny* or *Taf1d* signal (CB with 0 signal), as well as CBs with 1 or more than one signal, based on analysis of 170 CBs for *Ccny* and 225 CBs for *Taf1d*. H) RT-qPCR of isolated CBs (CB), or CB-depleted cytoplasmic fraction (Cytoplasm) after the treatment of seminiferous tubule segments with DMSO or STC-15 using *Taf1d* and *Ccny* primers. *Taf1d* and *Ccny* transcripts were quantified and normalized to input RNA. Data represent mean ± S.D. from four independent experiments; *P < 0.05. I) Western blotting with anti-DDX4 antibody to validate the equal CB isolation from DMSO and STC-15 treated cells. One representative IP is shown for each condition. SUP: CB-depleted cytoplasmic fraction.

The localization of SMG6 and PIWIL1 is strictly associated with the CB, and therefore, to address whether m⁶A-modified RNAs localize to the CB, we performed IF on mouse testis sections with an anti-m⁶A antibody. We showed that m⁶A signal is enriched in the nucleus but it also co-localizes with DDX4 to the CB from the precursor phase to mature CBs of round spermatids (Fig. 6C, Supplementary Figure S4). We also isolated CBs from mouse testes using our well-established protocol ^18^, and performed dot blotting of CB RNA with an anti-m⁶A antibody, which confirmed the enrichment of m⁶A-modified RNAs in the CB (Fig. 6D). Furthermore, m⁶A signal in the CB was reduced when germ cells isolated from adult testes were treated with an m⁶A inhibitor STC-15 for 24 hours before the CB isolation (Fig. 6E). Together, these data support the role for m⁶A modification in guiding the spatial localization of transcripts to specialized RNA granules for post-transcriptional regulation during spermatogenesis.

### m⁶A methylation regulates the CB localization of *Ccny* and *Taf1d* mRNAs

To explore the role of m⁶A modification in the SMG6/PIWIL1-mediated regulation of the selected target mRNAs *Ccny* and *Taf1d*, we first validated that these mRNAs are m⁶A-modified in the testis (Fig. 6A), which is consistent with a previous report that identified them as m⁶A-modified transcripts in differentiating male germ cells in mice ^35^. To study the effects of m⁶A modification on *Ccny* and *Taf1d* localization, segments of mouse seminiferous tubules from stages I-V of the seminiferous epithelial cycle ^3^ were cultured *ex vivo* and treated for 24 hours with either the m⁶A inhibitor STC-15, the m⁶A activator METTL3a1, or DMSO as a control. We then performed fluorescence *in situ* hybridization with probes detecting *Ccny* and *Taf1d* mRNAs combined with IF with DDX4 to visualize the CB (Fig. 6F), and quantified the *Ccny* and *Taf1d* mRNA signals overlapping with the CB (Fig. 6G). Quantification of CBs containing no RNA signal, one RNA signal, or more than one RNA signals, revealed that inhibition of m⁶A methylation reduced the localization of both transcripts to the CB, whereas activation of methylation with METTL3a1 increased their accumulation within CBs (Fig. 6G).

We further validated the effects of m⁶A inhibition on the CB localization of *Ccny* and *Taf1d* by RT-qPCR of CBs isolated from non-treated vs. STC-15-treated germ cells (Fig. 6H,I). We also collected a sample during the CB isolation protocol that represented the CB-depleted part of the cytoplasm ^18^. In the DMSO-treated control germ cells, *Ccny* and *Taf1d* were found in the CB fraction, indicating baseline processing under normal m⁶A conditions. Upon STC-15 treatment, the *Ccny* and *Taf1d* signals in the CB-depleted cytoplasm increased by approximately 40–60%, whereas CB enrichment decreased by roughly 40-50% relative to DMSO (Fig. 6H). These results indicate that m⁶A inhibition shifts these transcripts from CB-associated pools into the cytoplasmic fraction, which is consistent with the known role of m⁶A in promoting phase-separation of mRNAs and formation of RNP granules ^36^. Together, these data support a model in which m⁶A modification contributes to the spatial localization of mRNAs to the CB to undergo post-transcriptional regulation required for the progress of haploid gene expression program.

### Target mRNAs accumulate in the CBs of *Smg6-*deleted germ cells

To further investigate the interplay of SMG6 and PIWIL1 in mRNA regulation in the CB-environment *in vivo*, we analysed the levels of PIWIL1-associated *Ccny* and *Taf1d* mRNAs in germ cell-specific *Smg6* knockout (*Smg6*-cKO) testes compared with those in wild-type control testes via RIP followed by qPCR. Notably, we showed substantial increase in PIWIL1-bound *Ccny* and *Taf1d* mRNAs in *Smg6*-cKO testes compared to the control testes (Fig. 7A,B). This suggests that the CB-localized SMG6 acts downstream of PIWIL1, facilitating the clearance of PIWIL1-associated transcripts.

**Figure 7.**
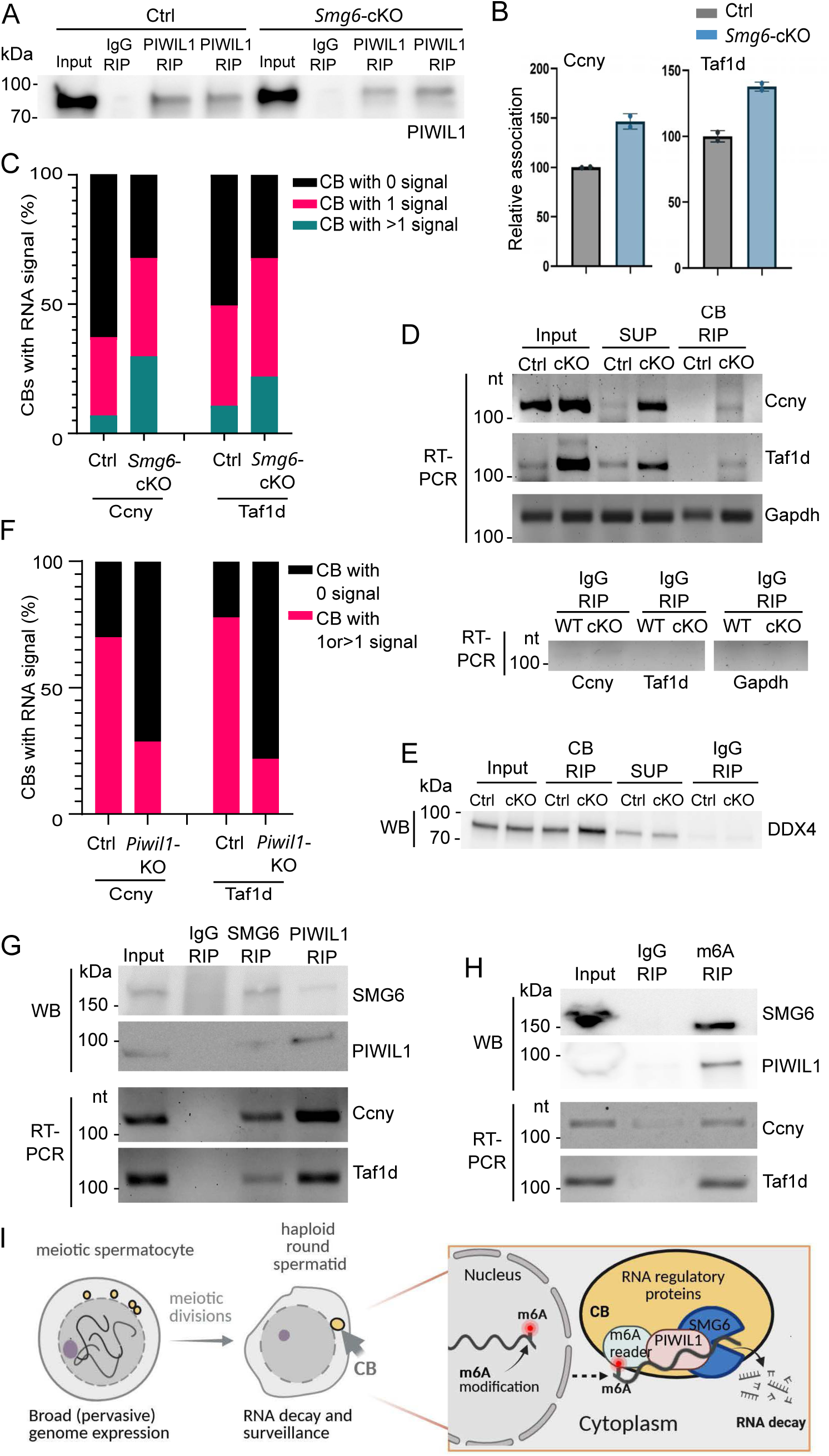
CB is involved in the SMG6-mediated regulation of *Ccny* and *Taf1d* mRNAs *in vivo*. A) Validation of the equal immunoprecipitation of PIWIL1 (PIWIL1 RIP) from control (Ctrl) and *Smg6*-cKO testes. Rabbit IgG was used as a negative control. B) anti-PIWIL1 RIP from control and *Smg6*-cKO testes followed by RT-qPCR with *Ccny* and *Taf1d* primers. *Taf1d* and *Ccny* transcripts were quantified and normalized to input RNA. Data represent mean ± S.D. from two independent experiments. D) Quantification of the *in situ* hybridization of *Ccny* and *Taf1d* mRNAs on control and *Smg6*-cKO testes sections (example images shown in Supplementary Fig. S5A). The graph shows the percentage of CBs with no overlapping *Ccny* or *Taf1d* signal (CB with 0 signal), as well as CBs with 1 or more than one signals, based on analysis of 151 CBs for *Ccny* and 185 CBs for *Taf1d*. D) RT-PCR of isolated CBs from control (WT) and *Smg6*-cKO (cKO) testes using primers for *Ccny, Taf1d* and *Gapdh*. Input: starting material for the IP. SUP: CB-depleted supernatant after low-speed pelleting of CBs. The lower panel shows the amplification in the negative control RIP with rabbit IgG. E) Validation of the successful CB isolation by Western blotting with anti-DDX4 antibody. The samples are as in (D). F) Quantification of the *in situ* hybridization of *Ccny* and *Taf1d* mRNAs on control and *Piwil1*-KO testes sections (example images shown in Supplementary Fig. S5B). The graph shows the percentage of CBs with no overlapping *Ccny* or *Taf1d* signal (CB with 0 signal), as well as CBs with 1 or more signals (CB with 1 or >1 signal), based on analysis of 150 CBs for *Ccny* and 119 CBs for *Taf1d.* G) SMG6 and PIWIL1 were immunoprecipitated from human testis lysate, followed by Western blotting with the same antibodies (upper panel) or RT-PCR with *CCNY* and *TAF1D* primers (lower panel). (H) Anti-m⁶A RIP from human testis lysate followed by Western blotting with anti-SMG6 and anti-PIWIL1 antibodies (upper panel) or RT-PCR with *CCNY* and *TAF1D* primers. I) Schematic diagram showing the appearance of the CB and summarizing our key findings on the CB-mediated RNA regulation (created with BioRender).

In line with the increased association of target mRNAs with PIWIL1 in the absence of SMG6, *Ccny* and *Taf1d* mRNAs were also shown to accumulate more in *Smg6*-cKO CBs than in control CBs (Fig. 7C-E). This was first visualized by *in situ* hybridization, which revealed a higher number of *Ccny* and *Taf1d* mRNA foci overlapping with the DDX4-positive CBs in *Smg6*-cKO round spermatids compared to the control (Supplementary Fig. S5A, Fig. 7C). Furthermore, to facilitate the quantification, we isolated CBs from *Smg6*-cKO and control testes, and showed elevated levels of intact *Ccny* and *Taf1d* transcripts in *Smg6*-cKO CBs relative to the control (Fig. 7D,E). The increased association of the target mRNAs with PIWIL1 and their accumulation in the CB in the absence of SMG6 indicate that the target recruitment to the CB and occurs normally, but transcript degradation is impaired in the absence of SMG6. Supporting the suggested role of PIWIL1/piRNAs in the recruitment of target mRNAs for CB-mediated regulation, *in situ* hybridization revealed that the CB-associated localization of *Ccny* and *Taf1d* was compromised in *Piwil1*-KO testes (Supplementary Fig. S5B, fig. 7F). These findings suggest that CBs serve as the primary site where PIWIL1-mediated target binding is coupled with SMG6 catalytic decay activity.

### SMG6-PIWIL1 association is conserved in human testis

Finally, to determine translational relevance, we examined whether the SMG6-PIWIL1 cooperation in the differentiating male germ cells is conserved in humans. The SMG6 and PIWIL1 sequences are well conserved between mice and humans (Supplementary Fig. S6). Immunoprecipitation from human testicular lysate confirmed that, similar to the mouse, SMG6 and PIWIL1 form complexes with each other and with *CCNY* and *TAF1D* mRNAs in the human testis (Fig. 7G). In addition, m⁶A RIP pulled down *CCNY* and *TAF1D* mRNAs, as well as SMG6 and PIWIL1 proteins from human testis lysate, demonstrating that the mRNA targets of the SMG6/PIWIL1 complex are also m⁶A modified in human (Fig. 7H). These findings support a role for the CB as a conserved hub for m⁶A-regulated and PIWIL1/SMG6-mediated RNA surveillance.

## DISCUSSION

In this study, we shed light on the molecular principles underlying CB-mediated RNA regulation in differentiating haploid male germ cells. The CB is an unusually large RNP granule, and its remarkable size likely reflects the exceptional demand for post-transcriptional RNA control during spermatogenesis ^21^. This demand arises from pervasive transcription during meiosis and the need to selectively stabilize, repress, or eliminate transcripts to enable the orderly transition from meiotic to the haploid gene expression program ^37^. Our findings reveal that the CB functions as an integrated regulatory hub that coordinates multiple RNA surveillance and regulatory pathways. Specifically, we demonstrate the convergence of PIWI–piRNA–mediated targeting, NMD endonuclease, and m⁶A RNA modification within the CB. While each of these pathways has independently been shown to safeguard male germline transcriptome integrity, our data uncover their functional cooperation to enable efficient and spatially organized post-transcriptional regulation essential for haploid male germ cell differentiation.

Our loss-of-function analyses support this cooperative logic. Depleting PIWIL1 or inhibiting m⁶A methylation markedly reduced SMG6 association with target transcripts and impaired their decay, and overexpressing SMG6 alone failed to rescue degradation in the absence of either cue. Conversely, depletion of SMG6 abolished piRNA-induced transcript decay, demonstrating that recognition by PIWIL1–piRNA is insufficient for clearance without an active endonuclease. Together, these observations uncover a three-way interdependence among m⁶A-dependent localization, piRNA-mediated targeting, and SMG6-driven catalysis.

Importantly, we recognise NMD endonuclease SMG6 as a novel effector protein in piRNA-mediated RNA regulation, thereby expanding the functional repertoire of the piRNA pathway. piRNAs have previously been shown to cooperate with CAF1-mediated deadenylation to promote elimination of mRNAs at the onset of spermatid elongation ^38^. Our findings demonstrate that instead of relying solely on deadenylation, piRNA-mediated target recognition can also engage the SMG6 endonucleolytic mechanism to achieve transcript decay. These results highlight a versatile post-transcriptional program in spermatids, in which piRNAs orchestrate large-scale mRNA clearance through multiple, complementary decay pathways.

The CB exhibits features of liquid–liquid phase separation (LLPS), analogous to stress granules and processing bodies that compartmentalize RNA metabolism via multivalent RNA–RBP interactions ^39^. In somatic cells, the m⁶A modification promotes selective partitioning of transcripts into condensates through YTH-domain and other m⁶A readers, enriching m⁶A-marked RNAs within stress granules and nuclear speckles ^25^. By analogy, our data propose that m⁶A functions as a spatial sorting signal that concentrates subsets of transcripts in the CB for regulated turnover. m⁶A reader-mediated bridging interactions could increase local encounter rates between m⁶A-tagged transcripts, PIWIL1– piRNA complexes, and SMG6, thereby facilitating efficient and selective decay. Notably, m⁶A recognition by YTHDF2 has been linked to UPF1-dependent NMD in somatic cells ^40^, revealing that this modification can serve as a permissive signal for engagement of the NMD machinery. UPF1 localizes to the CB together with SMG6 and m⁶A readers, supporting a potential role for UPF1 in CB-coordinated regulation of m⁶A-tagged transcripts.

While our findings substantially advance understanding of CB-mediated regulation, they also open exciting avenues for future mechanistic exploration. The open questions include the identity of the m⁶A readers responsible for SMG6 recruitment, the temporal relationship between m⁶A deposition and piRNA engagement, and the extent to which additional RNA decay pathways contribute to this process. Resolving these issues will be essential for a comprehensive understanding of how epitranscriptomic modifications and small-RNA-mediated regulation are integrated to sculpt the male germline transcriptome.

A central physiological outcome of the CB-centred RNA regulation is the control over the initiation and progression of the post-meiotic gene expression program. In the absence of SMG6 or PIWIL1, haploid cell differentiation fails to progress: spermatid elongation is not initiated, and germ cells are lost from the seminiferous epithelium ^17^. We propose that failure to clear specific transcripts at the correct time and place—due to loss of m⁶A-licensed CB entry, lack of piRNA-guided recognition, or absence of SMG6 catalysis prevents the transition from the meiotic to post-meiotic program, and the launch of the elongation program; this destabilizes cellular homeostasis, culminating in germ cell attrition. In this context, our candidate targets highlight how selective turnover can coordinate cell-cycle remodelling and transcriptional reprogramming: *Ccny* and *Taf1d*, identified in our degradome-seq and decay assays, are notable given their implicated roles in germ cell biology and transcriptional control ^29,30^.

In summary, our data support a model in which m⁶A licenses transcript recruitment to the CB, PIWIL1-piRNA complexes confer sequence specificity, and SMG6 mediates endonucleolytic cleavage (Fig. 7I). This coordinated division of labor enables precise and efficient clearance of transcripts required for spermatogenic progression. Disruption of any component—m⁶A deposition, CB integrity, or PIWIL1–SMG6 coupling may destabilize the germline transcriptome and contribute to male infertility. Importantly, here we demonstrate conservation of SMG6/PIWIL1 association in the human testis, and we have also confirmed CB-enriched SMG6 localization in human spermatids ^17^. Together with genetic links between piRNA and m⁶A pathway defects and human male infertility ^7,41^, these findings substantiate the critical role of these processes in the maintenance of human fertility. Therefore, we anticipate that CB-enriched RNA decay pathways may provide future opportunities for patient stratification and biomarker development in the diagnosis and management of male infertility.

## MATERIALS AND METHODS

### Animals

Mice were maintained and housed at the Central Animal Laboratory of the University of Turku, Finland, under controlled pathogen-free conditions in accordance with local regulations. Mice were euthanized by CO₂ inhalation followed by cervical dislocation. All animal experiments were approved by the Laboratory Animal Care and Use Committee of the University of Turku. The germline-specific conditional *Smg6* knockout mouse line has been previously described ^17^.

### Human samples

All persons included in the study gave written informed consent for the analysis of their donated material and the evaluation of their clinical data compliant with local requirements. The use of testicular tissue for pulldown analysis were approved by the Münster Ethics Committees/Institutional Review Boards (Ref. No. Münster: 2012-555-f-S)). Tissue samples were stored at −80 °C until processing.

### Isolation of germ cells

Round spermatids and spermatocytes were isolated from the testes of adult mice using a published protocol ^42^. Briefly, testes were decapsulated and digested sequentially with collagenase IV (Sigma, C5138) and DNase I (Sigma-Aldrich, DN25), followed by trypsin digestion (Worthington, LS003703) in 1× KREBS buffer. The resulting cell suspension was washed with 1× KREBS, filtered through a 100 µm cell strainer, and loaded onto a pre-chilled BSA gradient buffer. After 2 h of sedimentation at 4°C, germ cell fractions were collected, washed, and assessed for purity by DAPI staining.

### Co-immunoprecipitation (Co-IP) and RNA immunoprecipitation (RIP)

Mouse testes, human testicular biopsies, GC-2spd cells, or isolated mouse germ cells were homogenized in Pierce™ IP Lysis Buffer (Thermo Fisher Scientific, cat. no. 87787) supplemented with 1 mM DTT, 0.2 mM PMSF, RNase inhibitor (for RIP), and 1× protease inhibitor cocktail (Roche, cat. no. 11836170001). Homogenization was performed using a TissueLyser with 5 mm beads in a 96-well format (TissueLyser 5 mm Bead Dispenser, cat. no. 69975) and TissueLyser LT (cat. no. 85600) for 1 min at 50 Hz. Lysates were incubated on ice for 30 min and cleared by centrifugation at 14,000 × g for 10–20 min at 4°C. For Co-IP, lysates were pre-cleared with Protein G Dynabeads (Thermo Fisher Scientific, cat. no. 10003D) for 30 min at 4°C, and 100 µl of pre-cleared lysate was saved as an input. Antibody–bead complexes were prepared using 5 µg of SMG6, PIWIL1, or m⁶A antibodies, or control IgG (mouse or rabbit), and incubated at 4°C for 2 h. Lysates were then incubated with the antibody-conjugated beads overnight at 4°C. For RIP, the procedure was identical, except RNase inhibitors were included. Beads were washed three times with IP lysis buffer and resuspended in 100 µl lysis buffer. From this, 30 µl were used for Western blotting, and the remaining 70 µl were used for downstream RNA extraction and analysis.

### Western blotting (WB)

Protein samples were mixed with Laemmli buffer and denatured at 95°C for 5 min before sodium dodecyl sulfate–polyacrylamide gel electrophoresis (SDS-PAGE). Proteins were separated on 4–20% Mini-PROTEAN TGX precast gels (Bio-Rad, cat. no. 4561093) and transferred to PVDF membranes (Amersham, cat. no. RPN303F) using wet transfer. Membranes were blocked with 5% skimmed milk in 0.1% TBST for 1 h at room temperature and incubated overnight at 4°C with primary antibodies diluted in 5% skimmed milk, 0.1% Tween-20, 1× TBS. After three washes with 0.1% TBST, membranes were incubated with HRP-conjugated anti-rabbit or anti-mouse IgG (1:1000) for 1 h at room temperature. Proteins were detected using Western Lightning ECL Pro (PerkinElmer, cat. no. NEL120E001EA), imaged on a LAS4000 system (FujiFilm), saved as 16-bit TIFF files, and processed using ImageJ (NIH, version 1.8.0).

### Mass spectrometry

SMG6 and PIWIL1 Dynabead–antibody–protein complexes (3 replicates, prepared as described above) were washed three times with 50 mM Tris buffer (pH 8.0) at 4 °C and stored at −20 °C before analysis at the Turku Proteomics Facility. LC–ESI–MS/MS was performed using an Easy-nLC 1200 system coupled to a Q Exactive HF mass spectrometer (Thermo Fisher Scientific) with a nano-ESI source. Peptides were separated on a 15 cm C18 column (75 μm × 15 cm, ReproSil-Pur 5 μm 200 Å C18-AQ; Dr. Maisch HPLC GmbH) using a 40 min linear gradient of 8–43% acetonitrile (0.1% formic acid). Data were acquired with Xcalibur 4.1 and analyzed using Proteome Discoverer 2.4 (Thermo Fisher Scientific) with Mascot 2.7.0 (Matrix Science) against the SwissProt (2020) mouse database. The Fixed Value PSM Validator node was used for confidence scoring. SMG6- and PIWIL1-specific interacting proteins were defined as those detected exclusively in SMG6 or PIWIL1 co-IP samples, and only proteins identified by ≥2 unique peptides were retained. Over-representation analysis of Gene ontology (GO) terms, specifically Biological process (BP) and Molecular Function (MF), as well as Reactome pathways was performed using g:Profiler. Mus musculus was used as the reference organism. Statistical significance was determined using the default g:SCS thresholding method for multiple testing correction, with an adjusted p-value < 0.05 considered statistically significant.

### Isolation of chromatoid bodies

Chromatoid bodies (CBs) were isolated from adult mouse testes or *ex vivo*-cultured testicular cells following treatment, using a protocol adapted from ^43^ with minor modifications. Germ cells were released from testes of control and *Smg6*-cKO mice by incubation in 0.05% (w/v) collagenase (Worthington). The cell suspension was passed through a 100-µm strainer (BD Falcon), washed with phosphate-buffered saline (PBS), and either subjected to 24 hours culture or fixed immediately in 0.1% paraformaldehyde (Electron Microscopy Sciences) for 20 min at room temperature. Fixed cells were lysed by sonication (UCD-200, Diagenode) in 1 ml RIPA buffer (50 mM Tris-HCl, pH 7.5; 1% NP-40; 0.5% sodium deoxycholate; 0.05% SDS; 1 mM EDTA; 150 mM NaCl; supplemented with protease inhibitors (Roche), 0.2 mM PMSF, and 1 mM DTT) using six 30 sec pulses at medium power. The CB-enriched fraction was collected by centrifugation at 600 × g for 10 min, resuspended in RIPA buffer, and sonicated for two additional 30 sec intervals. Lysates were divided equally, and CBs were immunoprecipitated overnight at 4°C using Protein G Dynabeads (Thermo Fisher Scientific, cat. no. 10003D) coupled to anti-DDX4 antibody or rabbit IgG (negative control). For immunoblotting, aliquots of cross-linked cell lysate, CB IPs, IgG controls, and post-IP supernatants were mixed with Laemmli buffer, heated at 95°C for 5 min, and resolved by SDS–PAGE. For RNA studies, CB-associated RNA was processed as described below.

### m⁶A dot blotting

m⁶A levels were assessed by dot blotting using RNA isolated from SMG6 and PIWIL1 RNA IPs from adult mouse testis, CB–purified RNA fractions, and GC-2spd cells treated with DMSO or STC-15 (10 µM). For SMG6- and PIWIL1-RIP samples and CB RNA, the entire amount of purified RNA obtained from each preparation was applied to the membrane. For GC-2spd(ts) cells, 300 ng of total RNA was loaded per sample. RNA was denatured, applied to nylon membranes, and UV-crosslinked at 1200 µJ ×100 (5 min) using a UV crosslinker (insert instrument model). Membranes were blocked in 3% BSA in PBST and incubated with an m⁶A-specific antibody (Proteintech, Cat. No. 68055-1-Ig), followed by an HRP-conjugated mouse secondary antibody. Signals were detected by chemiluminescence imaging.

### piRNA extraction and detection

RNA was extracted from PIWIL1 or SMG6 RIP samples and corresponding IgG controls using a protocol similar to that described for RIP RNA extraction. RNA quality and size distribution were assessed using the Agilent Bioanalyzer. For small RNA analysis, isolated RNA was separated on 10-15% denaturing urea-polyacrylamide gels, post-stained with SYBR Gold (Invitrogen), and visualized using the ChemiDoc Imaging System (Bio-Rad).

### Transient transfection of PIWIL1-associated RNAs

PIWIL1-associated RNAs, including piRNAs, were isolated from PIWIL1 RIP samples as described above. Approximate concentration and RNA integrity were determined using an Agilent Bioanalyzer. GC-2spd cells were seeded in 6-well plates at 60–70% confluency one day prior to transfection. Transfections were performed using jetPRIME® DNA/siRNA (co-) transfection reagent (Polyplus-transfection, Cat No.: 101000001) according to the manufacturer’s instructions. Briefly, 40–60 ng of PIWIL1-associated RNA was diluted in jetPRIME® buffer and mixed with jetPRIME® reagent. After a 10-min incubation at room temperature, the RNA–reagent complexes were added dropwise to cells in complete medium. Cells were incubated for 24–48 h before harvesting. Control groups included IgG-associated RNAs and mock transfection (no RNA). Total RNA was then extracted from cells using TRIzol reagent (Invitrogen), and RT-PCR was performed to quantify mRNAs expression relative to *Gapdh* Expression levels were normalized and compared across experimental conditions.

### Immunofluorescence staining (IF)

Paraffin-embedded testis sections from adult *Smg6*-cKO, *Piwil1*-KO, and control mice were deparaffinized by sequential incubation: 3 × 5 min in xylene, 2 × 10 min in 100% ethanol, 2 × 10 min in 96% ethanol, and 2 × 10 min in 70% ethanol, followed by two washes in Milli-Q water (2 × 2 min). Antigen retrieval was performed in either sodium citrate buffer (10 mM sodium citrate, 0.05% Tween-20, pH 6.0) or Tris-EDTA buffer (10 mM Tris base, 1 mM EDTA, 0.05% Tween-20, pH 9.0) by boiling in a pressure cooker at 120°C for 20 min. Sections were blocked for 1 h at room temperature using either 10% normal donkey serum (Jackson ImmunoResearch, 017-000-121) and 3% BSA (Sigma, A2153) in PBST (0.1% Tween-20) or CAS-Block (Invitrogen, 00-8120). Primary antibodies were diluted (1:100–1:500) in blocking solution and incubated overnight at 4°C in a humidified chamber. The following day, sections were washed three times in PBST (5 min each) and incubated with Alexa Fluor 488/546/647-conjugated secondary antibodies (Life Technologies) at 1:500–1:1000 dilution for 1 h at room temperature. Slides were mounted using ProLong Diamond Antifade Mountant with DAPI (Life Technologies, P36962). Imaging was performed using a 3i CSU-W1 spinning disk confocal microscope (objectives: 40×, 63×, or 100×) with SlideBook6 software. Image processing was carried out using ImageJ (version 1.8.0, NIH).

### Antibodies

Primary antibodies used in this study were: DDX4 (ab13840), SMG6 (ab87539) PIWIL1 (ab12337), METTL3 (EPR18810) from Abcam; PIWIL1 (G82), normal rabbit IgG (2729S) from Cell Signaling Technology; hVasa (AF2030-SP) from R&D systems, m⁶A antibody (68055-1-Ig), PIWIL1 antibody (15659-1-AP), and mouse IgG (B900620) from Proteintech. Secondary antibodies conjugated with Alexa Fluor 488, 546, and 647 made in donkey were from Thermo Fisher Scientific (A-21202, A-21206, A-11055, A10036, A10040, A-11056, A-21203, A-21207, A-11058, A-31571, A-31573, A-21447). HRP-linked anti-rabbit IgG (NA934) and HRP-linked anti-mouse IgG (NA031) were from GE Healthcare Life Sciences.

### Cell Culture and Transfection

Mouse spermatogonial GC-2spd cells (ATCC® CRL-2196™, USA) were maintained in Dulbecco’s Modified Eagle Medium/Nutrient Mixture F-12 (DMEM/F-12, D-18900; Sigma-Aldrich) supplemented with 10% fetal bovine serum (FBS, S1810; Biowest) and 1% penicillin–streptomycin (15140-122; Gibco). Cultures were incubated at 37 °C in a humidified atmosphere containing 5% CO₂. For plasmid transient overexpression assays, GC-2spd cells were seeded in 6-well plates and transfected the following day with 2 µg of plasmid encoding for HA-SMG6 (a kind gift from Prof. Oliver Mühlemann) or FLAG-MIWI ^44^, or an empty pcdna3.1 (Invitrogen) as a control, at a 3:1 reagent-to-DNA ratio in Opti-MEM I (31985-062; Gibco). Cells were harvested 48 h post-transfection for downstream analyses. For siRNA or piRNA transfection including piRNAs isolated from PIWIL1 RIP samples, cells were transfected with 30–100 nM of siPOOLs, PIWIL1-associated RNAs, or synthetic piRNAs using jetPRIME® DNA/siRNA Transfection Reagent (101000001; Polyplus-transfection) following the manufacturer’s protocol. Briefly, RNA molecules were diluted in jetPRIME® buffer, complexed with jetPRIME® reagent for 10 min at room temperature, and added dropwise to cells in complete medium. All synthetic piRNAs used were 2′-O-methyl-modified at the 3′ end.

### Treatment of GC-2spd cells with m⁶A modulators

GC-2spd(ts) cells were cultured as described before. Following pre-culture, GC-2spd cells were treated with DMSO vehicle, the m⁶A inhibitor STC-15 (10 µM; HY-156677), or the METTL3 activator-1 (10 µM; HY-W037893) for 24–72 h. Treated cells were subsequently processed for downstream analyses, including RNA extraction, RNAscope *in situ* hybridization, and m⁶A dot blotting.

### *Ex-vivo* culture of mouse testicular cells and seminiferous tubules

Adult mouse testes were decapsulated, and testicular cells were released by enzymatic digestion with collagenase type IV (9001-12-1-C5138-100MG) for 15 min at 37 °C, passed through a 100 µm cell strainer, and collected by centrifugation at 600 × g for 5 min. Cells were plated in 24-well plates in DMEM/F12 medium (Sigma, D-18900) supplemented with 0.1% bovine serum albumin (BSA), 0.1% fetal bovine serum (FBS; Biowest, S1810), and 0.1% penicillin/streptomycin (Gibco, 15140-122). For seminiferous tubule culture, stage I–II or II–V tubules were microdissected ^45,46^ and placed in 20 µL of the same supplemented DMEM/F12 medium on glass slides within a humidified chamber. Both cell and tubule cultures were incubated at 34°C in 5% CO₂ for 24 h. For m⁶A studies, cells or tubules cultures were treated with m⁶A inhibitor STC-15 (10 µM; Cat. No.: HY-156677), m⁶A activator METTL3 activator-1 (10 µM; Cat. No.: HY-W037893), or vehicle control (DMSO; Sigma-Aldrich, Cat. No.: 472301). Following treatment, cells were subsequently processed for downstream analyses, including RNA extraction, RNAscope *in situ* hybridization, and m⁶A dot blotting, or western blotting to confirm CB isolation. For fluoresence *in situ* hybridization (FISH), testicular cells from stage I-V were spread from cultured seminiferous tubules into monolayers using the squash technique ^46,47^, snap-frozen in liquid nitrogen, fixed in acetone for 10 min, post-fixed in 4% paraformaldehyde, and subjected to RNAscope analysis targeting *Ccny* and *Taf1d*.

### Fluorescent *in situ* hybridization (FISH)

RNAscope® *in situ* hybridization was performed using the RNAscope Multiplex Fluorescent Reagent Kit v2 (Advanced Cell Diagnostics) following the manufacturer’s instructions. Three sample types were processed: paraffin-embedded testis cross-sections, squash-prepared seminiferous tubules, and GC-2spd(ts) cells. For paraffin-embedded testis sections, adult mouse testes were fixed, paraffin-embedded, and sectioned at 5 µm onto Superfrost Plus slides (Thermo Fisher Scientific). Sections were deparaffinized, treated with hydrogen peroxide, and subjected to heat-mediated target retrieval at 100 °C for 15 min, followed by protease digestion at 40 °C for 30 min. RNAscope probes targeting *Ccny* and *Taf1d* mRNAs were hybridized, and signal amplification and fluorophore development were performed according to the kit protocol. Nuclei were counterstained with DAPI and slides were mounted using ProLong™ Gold Antifade Mountant. Freshly prepared seminiferous tubule squashes and cultured GC-2spd(ts) cells were processed using the same RNAscope workflow with minor modifications. Prior to fixation, tubules or cells were treated for 24–72 h with either the m⁶A inhibitor STC-15 (10 µM; HY-156677), the m⁶A activator METTL3 activator-1 (10 µM; HY-W037893), or vehicle control (DMSO; Sigma-Aldrich 472301). Because these samples were not paraffin-embedded, heat-mediated antigen retrieval was omitted; instead, after fixation they underwent graded dehydration and rehydration before proceeding directly to protease treatment and subsequent hybridization.

### RNA extraction and reverse transcription

RNA was extracted from beads or input lysates using TRIzol reagent (Thermo Fisher Scientific) according to the manufacturer’s instructions and treated with DNase I (Sigma-Aldrich, AMPD1) to remove genomic DNA contamination. First-strand cDNA synthesis was performed using the Sensi-FAST™ cDNA Synthesis Kit (Bioline, cat. no. BIO-65053) following the manufacturer’s protocol.

### PCR and agarose gel electrophoresis

Target mRNAs were amplified from cDNA using gene-specific primers (Supplementary Table S4) under optimized PCR conditions on a Bio-Rad T100 Thermal Cycler. Reactions were performed using Thermo Fisher DreamTaq DNA Polymerase (5 U/µL, cat. no. EP0711) with 10× DreamTaq Green Buffer in a 50 µL reaction volume. PCR products were separated on 2% agarose gels containing Midori Green (Cat. No. MG04) and visualized using the FastGene FAS-DIGI PRO gel documentation system. Band intensities were quantified with ImageJ software (NIH, version 1.8.0).

### Quantitative PCR (qPCR)

cDNA was amplified using gene-specific primers (listed in Supplementary Materials) with Sensi-FAST™ SYBR® No-ROX Kit (Bioline, cat. no. BIO-98005) on a CFX96 Touch Real-Time PCR Detection System (Bio-Rad). Relative transcript levels in RIP samples versus input were calculated, and to assess changes in input samples under knockdown or overexpression conditions were normalized to housekeeping genes (Supplementary Table S4). Data was analyzed using GraphPad Prism 9.

### RNA pulldown assays using biotinylated DNA probes

Testis lysates were prepared in Pierce™ IP Lysis Buffer (Thermo Fisher Scientific, cat. no. 87787) as described above. 50 μL of Pierce™ Streptavidin Magnetic Beads (Cat. 88816) per sample were washed three times with wash buffer (TBS), resuspended in lysis buffer, and incubated with 100 pmol biotinylated DNA probes (control, *Taf1d*, or *Ccny*) for 90 min at room temperature with rotation. Beads were washed three times to remove unbound probe. Bead-bound probes were incubated with cleared lysate for 4 h at 4°C, followed by three washes with lysis buffer and a final wash with PBS. Bound proteins were eluted in SDS-PAGE sample buffer, heated at 95°C for 5 min, and analyzed by Western blot.

### RIP sequencing

Directional libraries without rRNA removal were prepared from three anti-SMG6 RNA immunoprecipitations (RIP), three anti-PIWIL1 RIPs, and corresponding IgG control RIPs, and sequenced on an Illumina NovaSeq X Plus Series using a paired-end 150-bp strategy at Novogene (UK) Company Limited. The quality of raw sequencing reads was assessed using FastQC (v0.11.9). Adapter sequences were removed and low-quality bases were trimmed using Trimmomatic (v0.39) ^48^. Clean reads were aligned to the mouse reference genome (Ensembl: Mus musculus GRCm38.101) using STAR (v2.7.10a) ^49^. Read assignment and quantification were performed with featureCounts (v2.0.3) ^50^. Raw read counts were normalized and differential expression analyses were carried out using DESeq2 (v1.44.0) ^51^. Publicly available transcriptome datasets representing distinct germ cell populations during mouse spermatogenesis (GSE35005) were obtained from the NCBI Gene Expression Omnibus^52^. Read mapping and normalization for the dataset was performed using the same pipeline as for the SMG6 and PIWIL1 RIP samples.

### Degradome sequencing

Three biological replicates of round spermatid samples from adult *Smg6*-cKO and control mice were submitted for degradome sequencing (degradome-seq, CD Genomics, Shirley, NY, USA). To construct degradome-seq libraries, polyA-RNA samples containing cleaved RNA fragments bearing 5′ monophosphates were ligated to RNA adaptor containing a 3’Mme I site and transcribed. After second-strand synthesis, MmeI digestion, gel purification, and PCR amplification, 3’ adaptor was ligated for deep sequencing with Illumina HiSeq SE50. After the quality check, reads were trimmed off adapters and low-quality bases were discarded using cutadapt (v.4.1). Reads were then aligned to the mouse genome (Ensembl: Grcm38) using HISAT2 (v.2.1.0) and assigned to the reference gtf file. Only sequences that originated from genes that were upregulated in *Smg6*-cKO round spermatids in RNA-seq ^17^ were kept, and they were further filtered according to their expression to keep only the ones with 10 or more counts in at least three individual samples among six samples (3 x control and 3 x cKO). Finally, differential expression analysis (based on unique sequences) was performed using DESeq2.

### Statistical analysis

Statistical comparisons between two groups were performed by Student’s t test using in R v4.1.3 and GraphPad Prism 9.0.0. Experimental replicates were performed as indicated in the respective Figure legends. The investigators were not blinded to allocation during experiments and outcome assessment.

## ACKNOWLEDGEMENTS

We thank all Kotaja lab members for their support and help. The Turku BioScience Cell Imaging Core Facility and Proteomics Core Facility, as well as Turku Center for Disease Modeling, TCDM, Turku, Finland, (www.tcdm.fi), are acknowledged for their help and services. Turku Central Animal Facility is acknowledged for providing facilities for animal maintenance and experimentation.

## AUTHOR CONTRIBUTIONS

A.A. conceived the study, performed experiments, analyzed the data, and wrote the manuscript; E.S. performed experiment and wrote the manuscript; L.M. and S.C. performed bioinformatics analyses and wrote the manuscript; A.D.P., E.V., S.O.M.L., R.P., R.A.G., K.T. and T.L. performed experiments; A.-K.D., S.K., N.N., B.S. and F.T. provided and handled human samples; J.-A.M. supervised the study, contributed to study design; N.K. conceived the study, supervised the study, contributed to the study design and wrote the manuscript. All authors contributed to finalizing the manuscript.

