## SUPPLEMENTARY INFORMATION for "Chromatoid body integrates piRNA, SMG6 and m⁶A pathways to control mRNAs in the male germline"

**Supplementary Figures S1-S6**

**Supplementary Table S1**: Mass spectrometric analysis of anti-SMG6 and anti-PIWIL1 IPs from adult mouse testes. A) SMG6-interacting proteins. B) PIWIL1-interacting proteins. C) Functional enrichment analysis of shared SMG6/PIWIL1-interacing proteins.

**Supplementary Table S2.** RIP-seq from adult testes with anti-SMG6 and anti-PIWIL1 antibodies. A) SMG6-associated mRNAs. B) PIWIL1-associated mRNAs. C) Functional enrichment analysis of shared SMG6/PIWIL1-bound mRNAs.

**Supplementary Table S3.** Degradome-seq analysis. A) List of shared upregulated genes (Log2FC > 1, Padj < 0.05) in *Smg6*-cKO and *Piwil1*-KO round spermatids. B) DE analysis of the degradation products identified in degradome-seq of *Smg6*-cKO vs. control round spermatids.

**Supplementary Table S4.** Primers and probes used in the study.

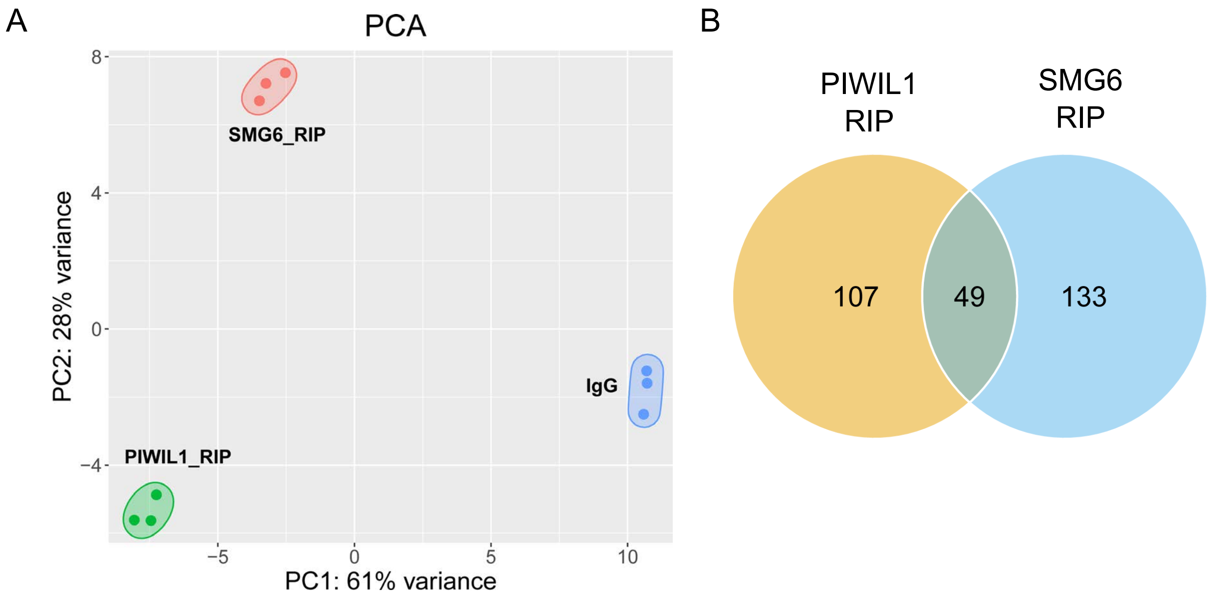


**Supplementary Figure S1.** A) Principal component analysis (PCA) plot shows the clustering of anti-SMG6, anti -PIWIL1 and IgG RIP-seq samples. Each point represents one biological replicate; ellipses indicate 95% confidence intervals. B) Venn diagram shows the overlap between SMG6- and PIWIL1-associated mRNAs identified by RIP-seq from adult mouse testes.

Supplementary Figure S2

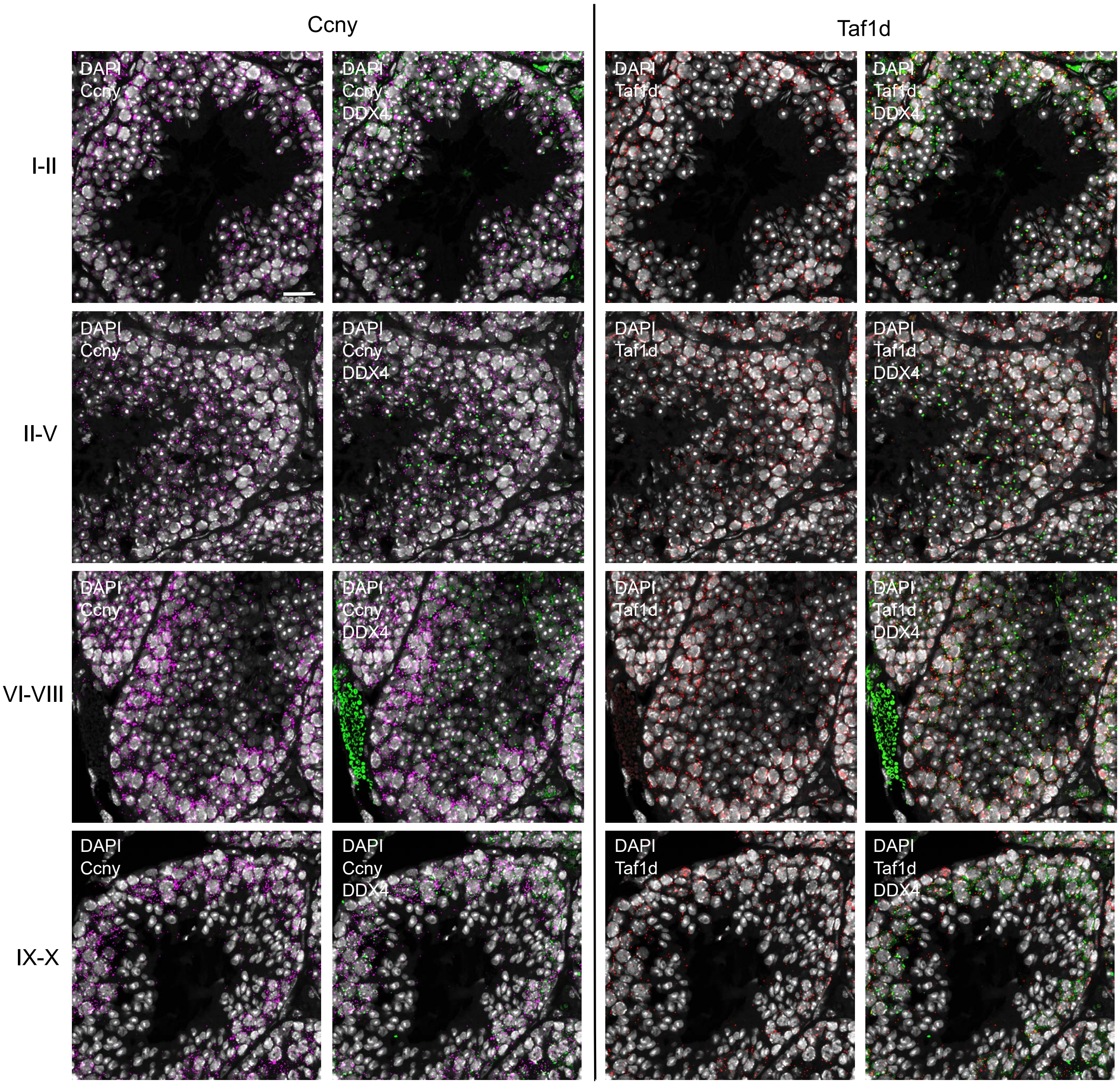


**Supplementary Figure S2.** *In situ* hybridization of PFA-fixed paraffin-embedded adult testis sections using specific probes detecting *Ccny* (magenta) and *Taf1d* (red). CBs are labelled using anti-DDX4 antibody (green). DAPI stains the nuclei (grey). Representative images of four stage ranges are shown (I, II-V, VII -VIII, IX-XI). Scale bar: 20 µm.

Supplementary Figure S3

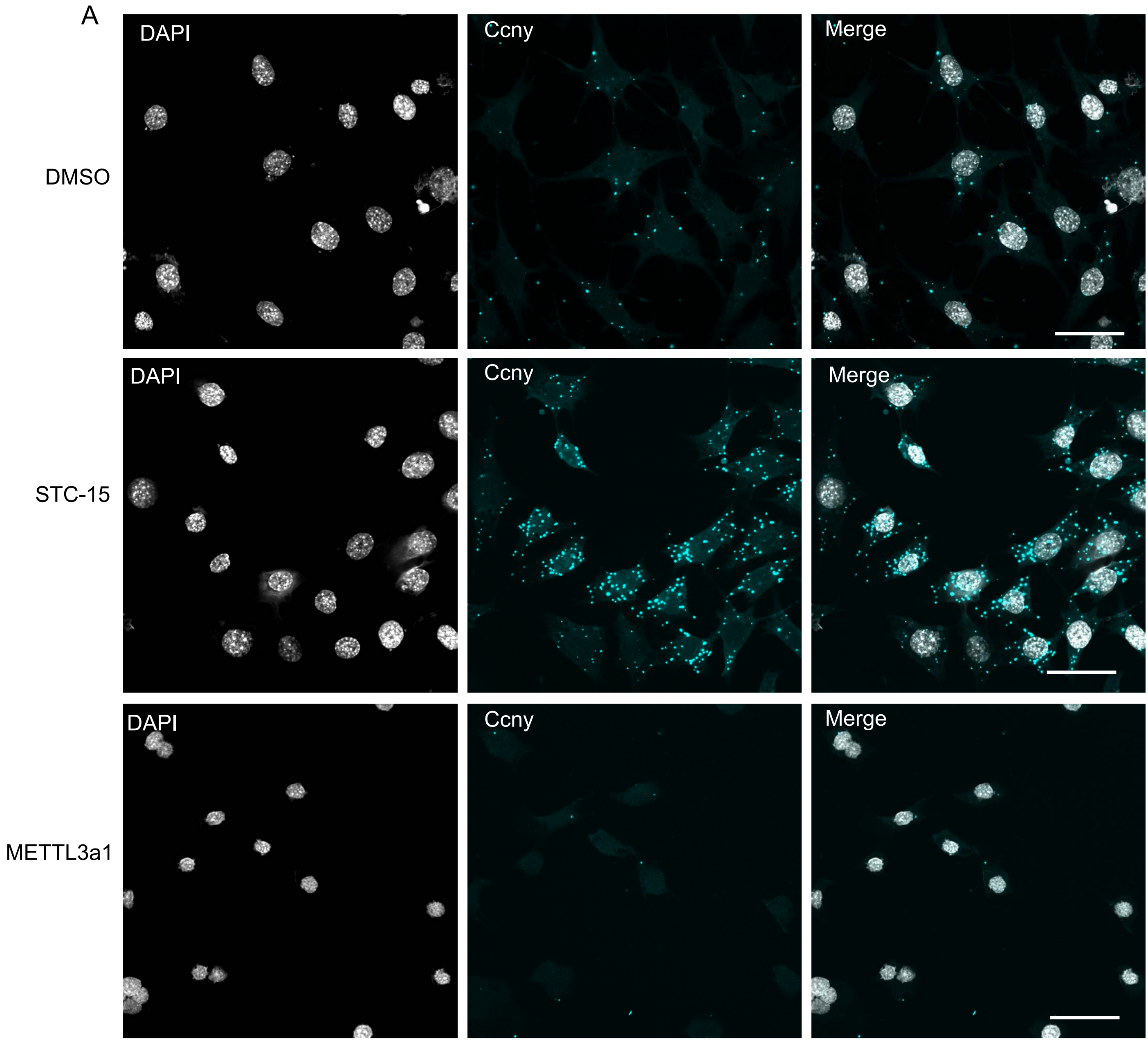


B

```
Ccny 3’UTR(-): 5' GTGCTGTCCCTCAGTTTCTCCTTTTGTT 3'
                           ||||||||||||||||||
piRNA:         3' UUUCGUUCAGAGUCAAAGAGGAAAACAU 5'
```

**Supplementary Figure S3.** A) *In situ* hybridization of cultured GC-2spd cells with *Ccny*-specific probe (cyan) after treatment of cells with DMSO (control), STC-15 or METTL3a1. DAPI stains the nuclei (grey). Scale bar: 10 µm. B) Pairing of *Ccny*-targeting piRNA with *Ccny* mRNA.

Supplementary Figure S4

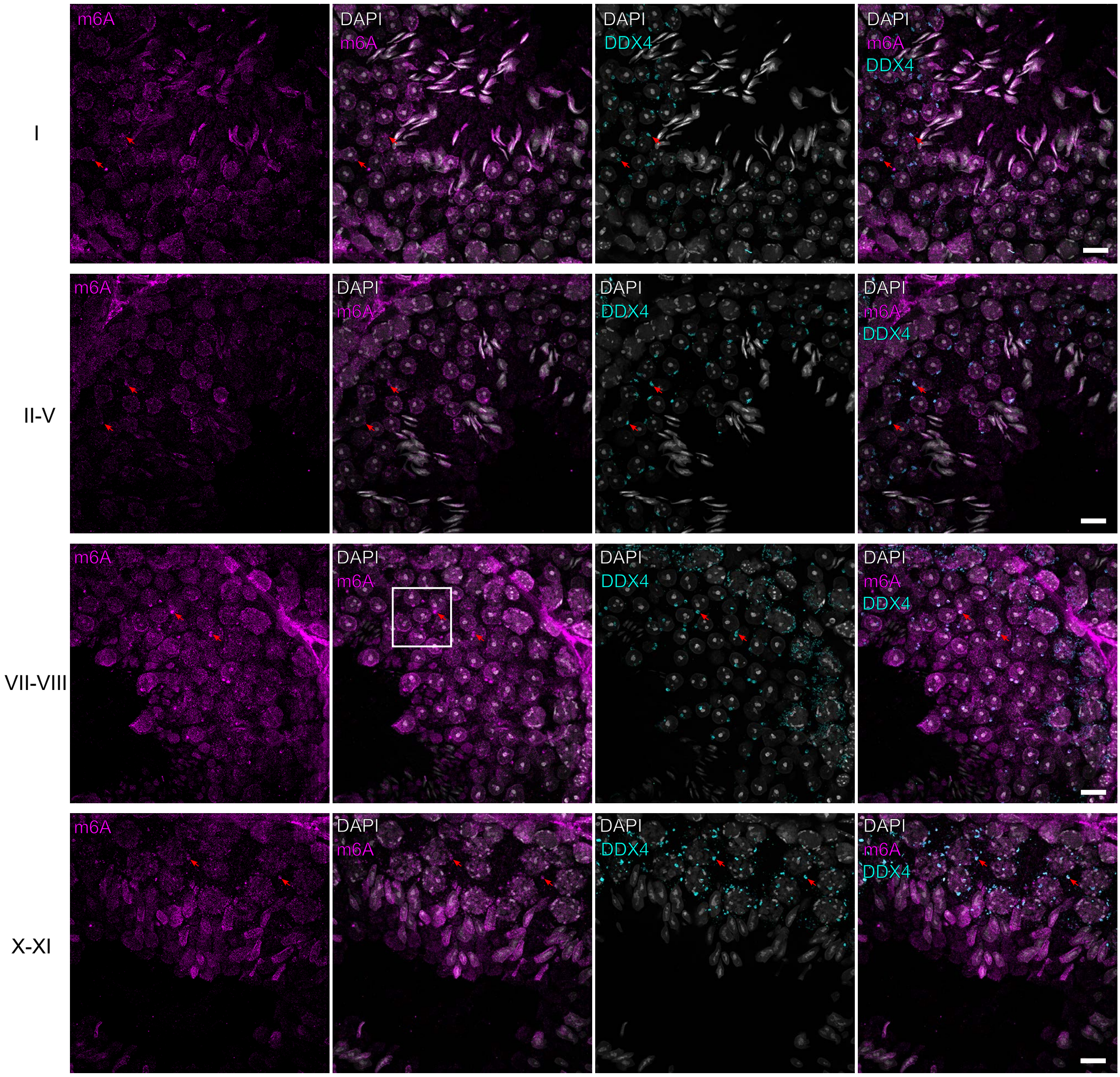


**Supplementary Figure S4.** IF staining of PFA -fixed paraffin-embedded adult testis sections using anti-m⁶A antibody (magenta). DDX4 antibody was used to label the CBs (cyan). DAPI stains the nuclei (grey). Representative images of the stages I, II-V, VII-VIII and -XI of the seminiferous epithelial cycle are shown. Some examples of the CBs (stages I, II-V, VII-VIII) or CB precursors (stage X-XI) are pointed by red arrows. The area shown in Figure 6C is indicated with a white box. Scale bar: 10 µm.

Supplementary Figure S5

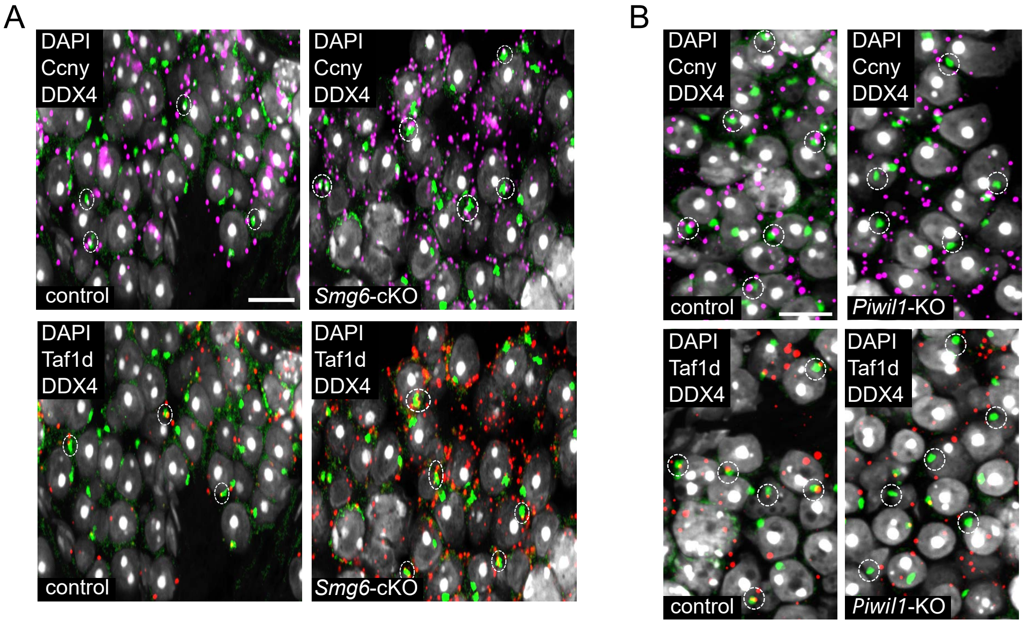


**Supplementary Figure S5.** A) *In situ* hybridization of *Ccny* (magenta) and *Taf1d* (red) mRNAs on control and *Smg6*-cKO testes sections. CBs were labeled with anti -DDX4 (green). DAPI stains the nuclei (grey). Scale bar: 10 µm. B) *In situ* hybridization of *Ccny* (magenta) and *Taf1d* (red) mRNAs on control and *Piwil1*-KO testes sections. CBs were labeled with anti-DDX4 (green). DAPI stains the nuclei (grey). Scale bar: 10 µm.

Supplementary Figure S6

*SMG6*

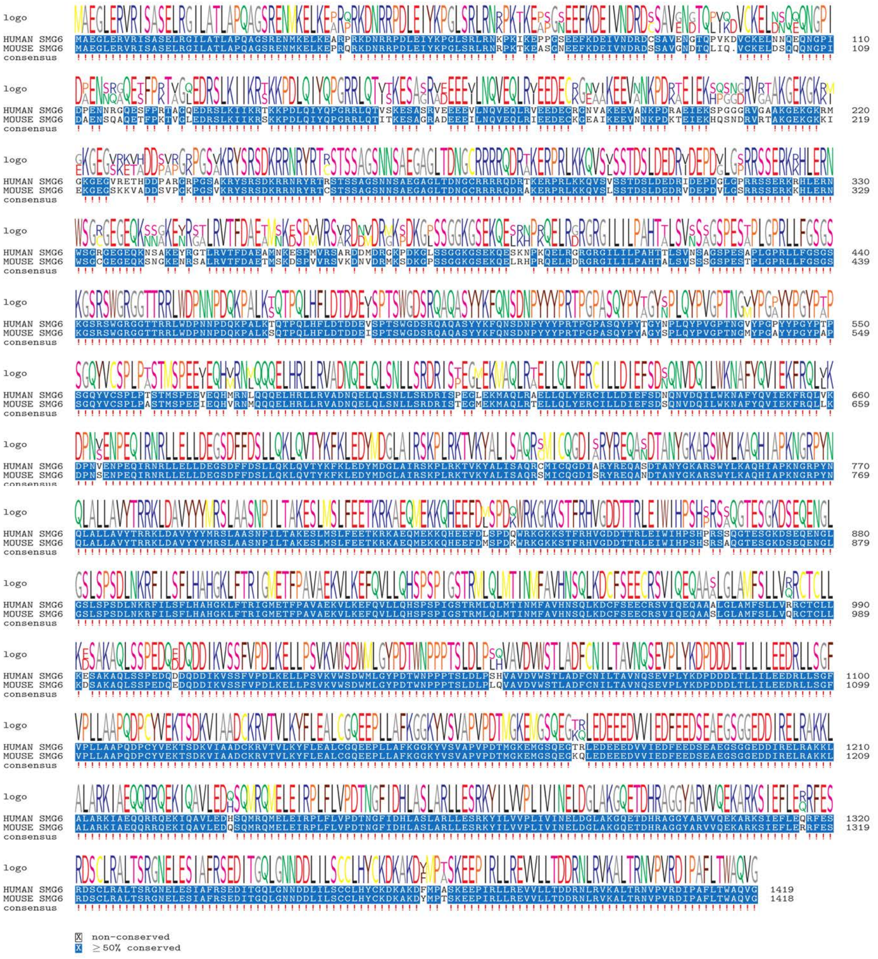


*PIWIL1*

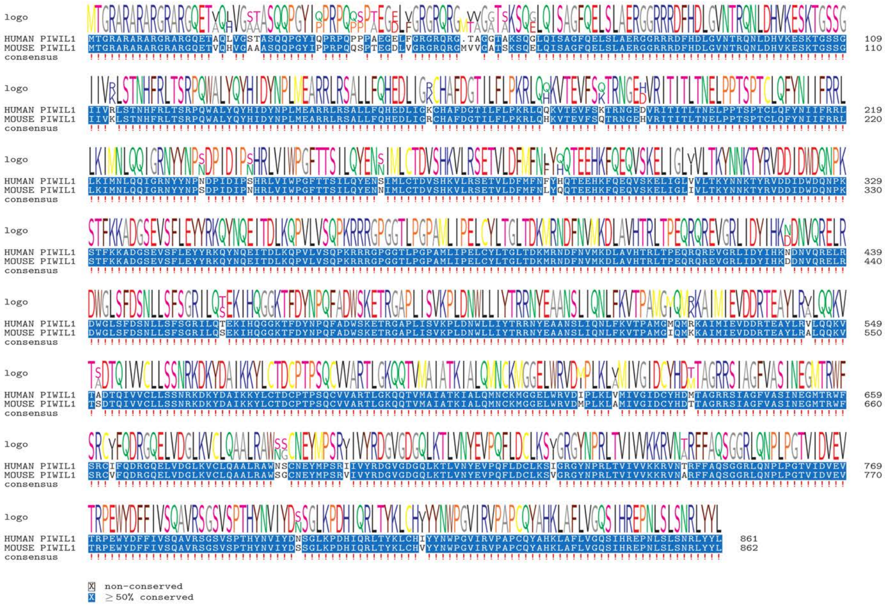


**Supplementary Figure S6.** Pairwise protein sequence alignments of human and mouse SMG6 and PIWIL1. Orthologous protein sequences from Homo sapiens and Mus musculus were aligned to assess conservation. Sequences are identified by UniProtKB accessions: Q86US8 (human SMG6) compared wi th P61406 (mouse SMG6), and Q96J94 (human PIWIL1) compared with Q9JMB7 (mouse PIWIL1). Blue highlights indicate positions of amino acid identity between the species pairs.
